# Neutrophils mediate the capture of peritoneal contaminants by fat-associated lymphoid clusters of the omentum

**DOI:** 10.1101/774968

**Authors:** Lucy H. Jackson-Jones, Peter Smith, Marlène S. Magalhaes, Jordan R. Portman, Katie J. Mylonas, Mark Nixon, Beth E.P. Henderson, Ross Dobie, Neil C. Henderson, Damian J. Mole, Cécile Bénézech

## Abstract

The omentum is a visceral adipose tissue rich in fat-associated lymphoid clusters (FALCs), which collects peritoneal contaminants and provides a first layer of immunological defence within the abdomen. Using single-cell RNA sequencing and spatial analysis of omental stromal cells, we reveal that the surface of FALCs are covered with specialised mesothelial cells, which we name FALC cover cells. We demonstrate that CXCL1 is expressed by FALC cover cells and that CXCL1 is critical for the retention and accumulation of neutrophils within FALCs during peritonitis. We show that protein arginine deiminase 4 mediates the formation of dense neutrophil aggregates, which are required for the neutralisation of particles present in the peritoneal cavity. Finally, we provide confirmatory evidence in humans with acute appendicitis, that the omentum is also a site of neutrophil recruitment and bacterial capture, and is thus an important component of the immunological defence against the propagation of peritoneal contaminants.

## Introduction

The omentum, a visceral fat depot contained within a fold of peritoneum, has the capacity to rapidly absorb particles and pathogens present in the peritoneal cavity (Meza-Perez and Randall, 2017; Platell et al., 2000). The omentum is an immune-adipose organ, the immunological properties of which are derived from the presence of numerous immune cell clusters called fat-associated lymphoid clusters (omFALCs, historically named milky spots) (Rangel-Moreno et al., 2009) which are also found in the mesentery (Moro et al., 2010), the mediastinum and pericardium, in association with the peritoneal, pleural and pericardial cavities (Benezech et al., 2015; Jackson-Jones et al., 2016). The continuous flow of fluid from the peritoneal cavity through omFALCs makes them unique niches for the clearance of peritoneal contaminants and initiation of protective immune responses during peritonitis. Peritonitis remains the second leading cause of sepsis and death in patients in intensive care units (Montravers et al., 2016; Ross et al., 2018).

FALCs support multifaceted stromal-immune cell interactions which are critical for the maintenance and function of innate-like B cells within the serous cavities as well as facilitating T-cell dependent B cell immune responses to peritoneal antigens (Ansel et al., 2002; Benezech et al., 2015; Rangel-Moreno et al., 2009). FALC stromal cell derived CXCL13 maintains peritoneal innate like B Cells (Ansel et al., 2002; Benezech et al., 2015; Rangel-Moreno et al., 2009). Upon inflammatory signals, serous B cells migrate into FALCs where the provision of IL-5 by type 2 innate lymphocyte (ILC2) enables rapid B cell proliferation and IgM secretion(Jackson-Jones and Benezech, 2018; Jackson-Jones et al., 2016). This is dependent on IL-33, produced by FALC stromal cells (Jackson-Jones et al., 2016) which induces the production of IL-5 by ILC2 (Moro et al., 2010). Peritonitis induces *de novo* FALC formation, which is dependent on the production of TNF by monocyte/macrophages and TNFR-signalling in stromal cells (Benezech et al., 2015). The initial recruitment of inflammatory monocytes into FALCs is dependent on MYD88 dependent activation of *Ccl19* expressing FALC stromal cells (Perez-Shibayama et al., 2018). The cross-talk between monocyte/macrophage and FALC stromal cells is required to support B cell differentiation (Perez-Shibayama et al., 2018) and FALC expansion (Benezech et al., 2015).

FALCs are highly vascularised (Cruz-Migoni and Caamano, 2016; Meza-Perez and Randall, 2017) and act as gateways to the peritoneal cavity during peritonitis. Work by Buscher et al., demonstrated that during TNFα induced peritonitis, neutrophils very rapidly transit through the high endothelial venules of omFALCs to enter the peritoneal cavity (Buscher et al., 2016). The extravasation of cells into the peritoneal cavity via FALCs may be facilitated by the presence of a loose lining of mesothelial cells (Doherty et al., 1995; Hodel, 1970).

The ability of the omentum to collect peritoneal contaminants is widely attributed to the flow of fluid from the peritoneal cavity through the omentum with omFALCs acting as an integrated filtration system. However, little is known about how FALCs perform this function and whether stromal-immune cell interactions active within omFALCs contribute to this phenomenon. Using single-cell RNA sequencing (scRNAseq), we reveal that the stromal compartment of the murine omentum is heterogenous and supports an elaborate three-dimensional organisation of omFALCs. In particular, we define two populations of mesothelial derived stromal cells covering the surface of FALCs that are specialised in the production of immune mediators including the neutrophil recruitment chemokine CXCL1. We show that CXCL1 is critical for the retention and accumulation of neutrophils in omFALCs in response to Zymosan-induced peritonitis. At these sites, neutrophils form dense aggregates that are coated with neutrophil extra-cellular trap (NET)-like DNA structures and which concentrate Zymosan particles. *In vivo* chemical inhibition of protein arginine deiminase 4 (PAD4) completely abolishes omental-neutrophil aggregation, preventing the capture of Zymosan particles by FALCs resulting in increased dissemination of peritoneal contaminants to the spleen. Finally, we provide evidence that similar NET-like DNA structures within the omentum are involved in the collection of bacterial contaminants during human peritonitis.

## Results

### scRNAseq reveals the presence of three distinct FALC stromal cell populations

To characterise the mesothelial & stromal cell populations of the omentum we performed droplet-based scRNAseq on isolated mouse omental CD45^−^CD41^−^Ter119^−^CD31^−^ stromal cells (Fig. 1A) from naïve mice. To identify distinct cell populations based on shared and unique patterns of gene expression, we performed unsupervised clustering using the Seurat software package, and identified five populations which we visualised using t-SNE and a hierarchical cluster tree (Fig. 1B-C). We identified differentially expressed genes (DEG; genes with a 0.25 log-fold change and expressed in at least 25% of the cells in the cluster under comparison) for each cluster. We identified the largest cluster as mesothelial cells (Fig. 1B, red) as DEGs were enriched for epithelial (*Upk1b*, *Upk3b*, *Krt19*, *Krt7*) and mesothelial (*Msln* and *Cd200)* lineage marker genes (Fig. 1D,E,F and Supplementary Fig. 1A). Two clusters (Fig. 1B,C pale and dark green) shared similarity with mesothelial cells in keeping with the expression of some epithelial lineage marker genes such as *Upk1b*, *Upk3b* and *Krt19* (Fig. 1F and Supplementary Fig. 1A). Remarkably, one of the two additional mesothelial like clusters (pale green, *Cxcl13^+^* mesothelium) was distinguished by DEG involved in the recruitment, adhesion or activation of immune cells such as *Hdc*, *Enpp2*, *Ccl2*, *Cd44, Il34*, *Cxcl10*, *Cxcl13*, *Cd55*, *Ctsc*, *Ccl7* and *Cxcl1* (Fig. 1D,E,G, Fig. 2D and Supplementary Fig. 1B). The existence of a population of CXCL13^+^ stromal cells localised around the outside of FALCs has previously been reported (Benezech et al., 2015; Rangel-Moreno et al., 2009). The fact that these cells expressed mesothelial markers suggested that *Cxcl13^+^* cells were covering the surface of FALCs. The second mesothelial like cluster (dark green, *Ifit^+^* mesothelium) was distinguished by DEGs associated with interferon signalling such as *Ifit3b*, *Ifit3*, *Ifit1* and *Isg15* and anti-viral responses such as *Rsad2* (Fig. 1D,E,H and Supplementary Fig. 1C). Pathway analysis confirmed association of this cluster with interferon signalling and anti-viral mechanism terms (Supplementary Fig. 2). This suggested that cells from these two clusters derived from mesothelial cells and acquired specific immune functions.

**Figure 1:**
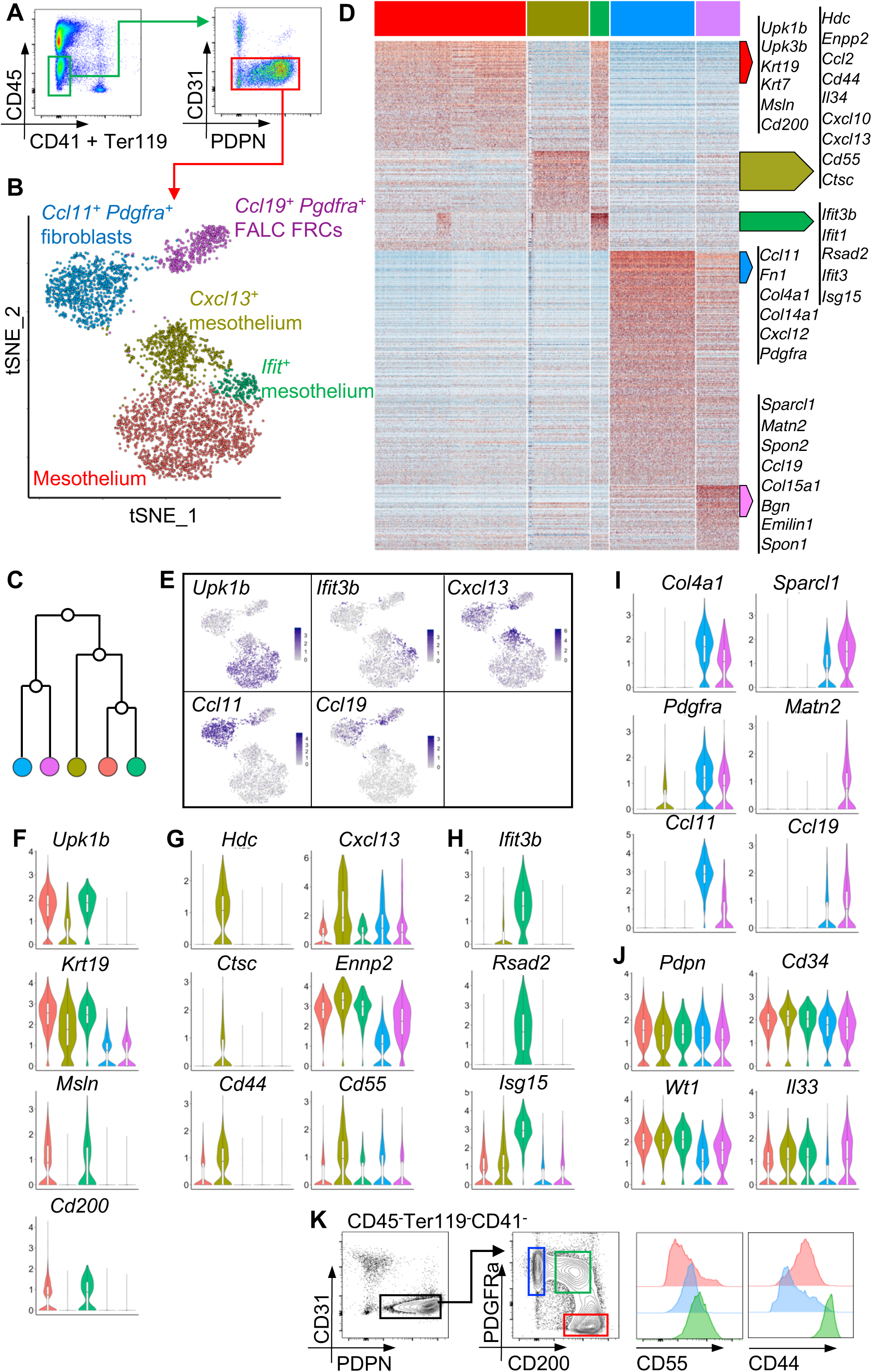
Identification of five non-endothelial stromal cell populations within the omentum. **A** CD45^−^CD41^−^Ter119^−^CD31^−^ non endothelial omental stromal cell were cell-sorted and underwent droplet-based scRNAseq. **B-C,** Unsupervised clustering of non-endothelial omental stromal cells visualized with tSNE where each dot is a single cell colored by cluster assignment (**B**) and a hierarchical cluster tree (**C)**. **D,** Heatmap of each cell’s (column) expression of DEGs (row) per cluster. **E,** Gene expression distinguishing the five clusters projected onto tSNE plots. Colour scaled for each gene with highest log-normalized expression level noted. **F-J,** Violin plots of canonical omental stromal cell gene expression by cluster with highest log-normalized expression value labelled. **K,** Representative gating strategy of non-endothelial omental PDPN^+^PDFGRa^+^CD200^−^ (blue), PDPN^+^PDFGRa^int^CD200^int^ (green), and PDPN^+^PDFGRa^−^ CD200^+^ (red) stromal cells and level of expression of CD55 and CD44 in these populations.

**Figure 2:**
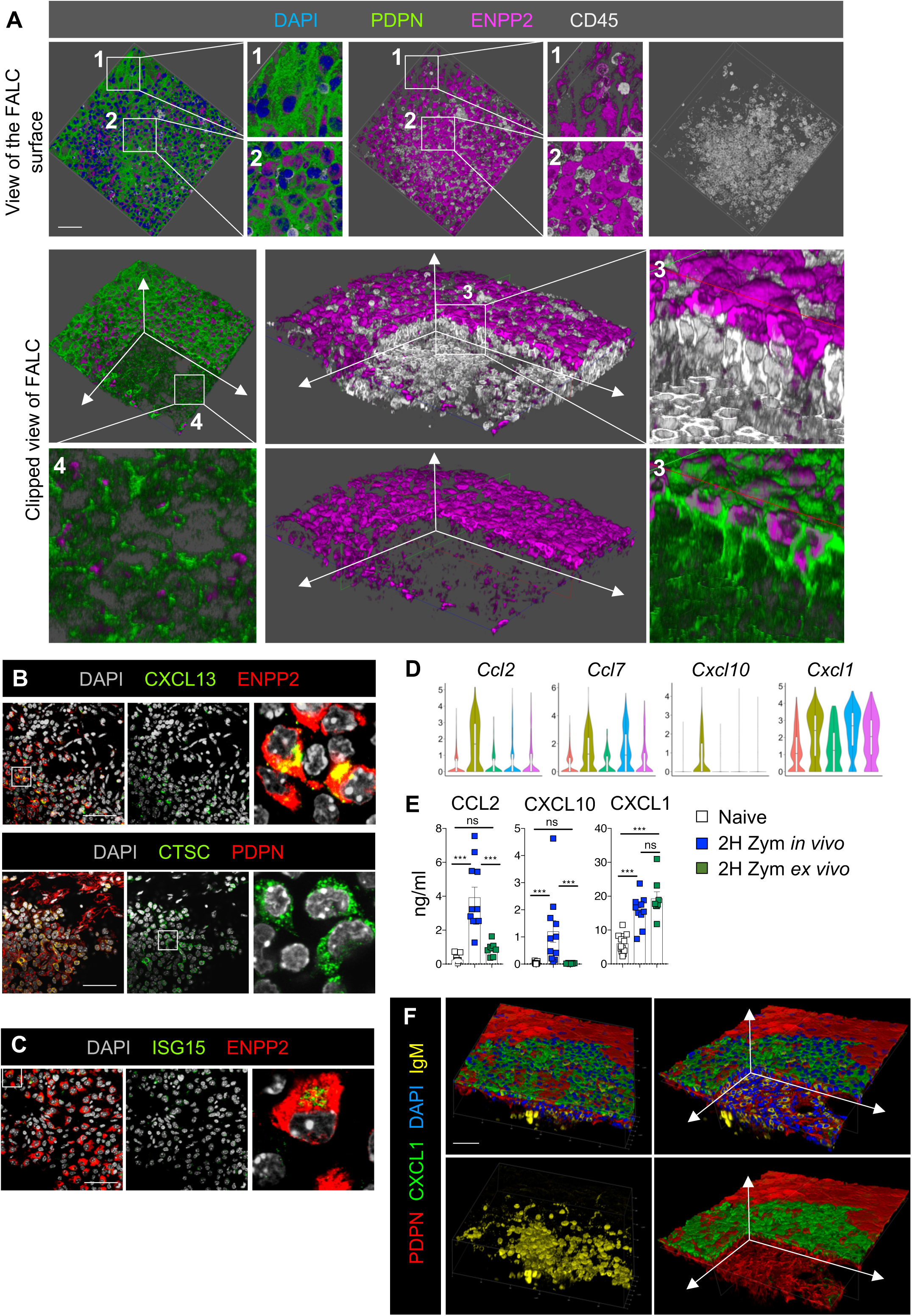
FALCs are covered by a differentiated monolayer of mesothelial cells. **A,** 3D reconstruction of omFALC obtained by confocal imaging of wholemount immunofluorescence staining of the omentum showing the surface of the cluster (first row) and two clipped views of the inside of the cluster (second row). Four enlargements are shown: 1 for mesothelial cells, 2 and 3 for FALC cover cells and 4 for FALC FRCs. **B-C,** Representative confocal images of wholemount immuno-fluorescence staining of omentum showing the surface of omFALCs and the expression of CXCL13, Cathepsin-C and ISG15 by FALC cover cells. **D,** Violin plots of the gene expression of the inflammatory chemokines *Ccl2, Ccl7, Cxcl10* and *Cxcl1* by cluster with highest log-normalized expression value labelled. **E,** Amounts of CCL2, CXCL10 and CXCL1 secreted into the supernatant of 2h omentum explant culture per omentum and per ml after exposure to Zymosan-A for 2h either *in vivo* (i.p. injection) or *ex vivo.* Data pooled from two independent experiments with n=10 mice per group. **F,** 3D reconstruction of omental FALC obtained by confocal imaging of wholemount immunofluorescence staining of the omentum showing the surface of omFALCs (first column) and a clipped view inside the cluster (second column). Scale bar 50 μm. All staining representative of n≥4 mice in at least 2 independent experiments. Error bars show SEM. Kruksal Wallis test with Dunn’s multiple comparisons test or ANOVA with Sidak’s multiple comparisons test were applied after assessing normality using D’Agostino & Pearson Normality test, ns= non-significant, *** P=<0.001.

The remaining two clusters (Fig. 1B, blue and purple), displayed DEGs enriched for genes associated with fibroblasts such as *Fn1*, *Col4a1*, *Col14a1* and *Pdgfra* (Fig. 1D,I and Supplementary Fig. 1D). The largest cluster (blue) was distinguished by the expression of *Ccl11* (also called Eotaxin) and assigned the name *Ccl11^+^Pdgfra^+^* fibroblasts (Fig. 1B,E,I). The final cluster of fibroblasts (purple) was characterised by the expression of *Ccl19*. The existence of a dense network of PDFGRα^+^ fibroblast reticular cells (FRC) expressing *Ccl19* and embedding FALC B cells was recently described in the omentum and mesenteries (Perez-Shibayama et al., 2018). This cluster showed enrichment for genes involved in formation of the extra-cellular matrix (ECM) characteristic of lymph node (LN) and FALC FRCs such *as Sparcl, Matn2, Spon2, Col15a1, Bgn, Emilin1* and *Spon1* (Fig. 1D,I and Supplementary Fig. 1E) (Huang et al., 2018; Malhotra et al., 2012; Perez-Shibayama et al., 2018). We thus assigned this cluster the name *Ccl19^+^ Pdgfra^+^* FALC FRCs. They represented a distinct subset of FRCs whose gene expression profile did not fit any of the LN stromal cell populations recently described by scRNAseq (Rodda et al., 2018). In particular FALC FRCs did not express *Il7*, *Ccl21*, *Bst1* or *Cxcl9* but expressed high levels of *Cd34* (Fig. 1J), *Inmt*, and *Nr4a1* (Supplementary Fig. 1E).

All five of the identified clusters expressed *Pdpn* and the genes *Wt1*, *Cd34*, *Itgb1* (*Cd29*) and *Ly6a* (*Sca-1*) which have been used collectively to identify cells of mesothelial origin giving rise to adipocytes in visceral fat depots (Fig. 1J and Supplementary Fig. 1F) (Chau et al., 2014). Regulation of retinol metabolism by *Wt1^+^* expressing cells was recently shown to be critical to maintain GATA6^+^ resident macrophages in the peritoneal cavity (Buechler et al., 2019). We confirmed expression of the 2 step-limiting enzymes of retinol metabolism by omental stromal cells with *Aldh1a1* highly expressed by mesothelial cells and FALC cover cells and *Aldh1a2* highly expressed by mesothelial cells and *Pdgfra^+^Ccl11^+^* fibroblasts (Supplementary Fig. 1F). Finally, we found that *Il33* was expressed by all three mesothelial clusters, in agreement with recent reports (Mahlakoiv et al., 2019; Spallanzani et al., 2019) as well as in FALC FRCs, confirming our previous observation that IL-33 was expressed by FALC stromal cells (Jackson-Jones et al., 2016).

The existence of PDPN^+^PDGFRα^+^ and PDPN^+^PDGFRα^−^ stromal cells in the mesenteric adipose tissue was recently described (Koga et al., 2018). To refine the flow-cytometric analysis of FALC rich tissues we utilised the cell surface markers CD55 and CD44 and confirmed the presence of PDGFRα^−^PDPN^+^CD200^+^CD55^−/low^CD44^low^ mesothelial cells and PDGFRα^+^PDPN^+^CD200^−^CD55^int^CD44^−^ fibroblasts in the omentum. In addition we could identify a population of cells transitioning from PDGFRα^−^CD200^high^ to PDGFRα^+^CD200^−^ and expressing high levels of CD55 and CD44 in keeping with the gene expression profile of *Cxcl13^+^* FALC stromal cells (Fig. 1K).

### FALCs are covered by a monolayer of stromal cells expressing high levels of ENPP2

To elucidate the spatial organisation of FALCs, we performed wholemount immuno-fluorescence staining using marker genes identified above. Staining for the lysophospholipase ENPP2 (Ectonucleotide Pyrophosphatase/ Phosphodiesterase 2), expression of which was enriched in the *Cxcl13^+^* FALC stromal cell population, revealed the existence of PDPN^+^ cells covering the surface of omFALCs and expressing very high levels of ENPP2. The morphology of these cells differed from the typical cobblestone appearance of PDPN^+^ mesothelial cells which surrounded the FALC and expressed low levels of ENPP2. PDPN^+^ FALC FRCs, which formed a reticular network at the core of the cluster, expressed very low levels of ENPP2 as predicted (Fig. 2A). We confirmed the presence of the B-cell positioning chemokine CXCL13 and of the lysosomal cysteine protease Cathepsin-C (*Ctsc*; also known as di-peptidyl peptidase I), within the stromal cells covering omFALCs (Fig. 2B). Our results thus confirmed the existence of a mesothelial derived population of cells covering the surface of FALCs and expressing ENPP2, CXCL13 and Cathepsin-C, which we named *Cxcl13^+^* FALC cover cells. We used ISG15 to track the presence of *Ifit^+^* FALC stromal cells and found the expression of ISG15 in the cytoplasm of a subset of the ENPP2^+^ cells covering omFALCs (Fig. 2C). We named these cells *Ifit^+^* FALC cover cells. Finally, staining for CCL11 revealed that fibroblasts contained in the adipose (non-FALC) stroma of the omentum expressed high levels of CCL11, while FALC FRCs and mesothelial cells did not (Suppl. Fig. 3).

### Cxcl13^+^ FALC cover cells express inflammatory chemokines

The *Cxcl13^+^* FALC cover cell cluster was distinguished by the expression of the monocyte chemo-attractants *Ccl2* and *Ccl7* and the neutrophil chemo-attractants *Cxcl1* and *Cxcl10* (Fig. 2D) suggesting a role for these cells in orchestrating the recruitment of inflammatory cells during peritonitis. The expression of *Cxcl1* was also enriched in *Ccl11^+^Pdgfra^+^* stromal cells and FALC FRCs. Analysis of omental explant culture supernatants showed that CXCL1 protein was released at steady state by the omentum and that CXCL1 secretion was potentiated after a two-hour exposure to Zymosan-A, *in* or ex *vivo* (Fig. 2E). In contrast, the two other early chemo-attractants, CCL2 and CXCL10, were found to be released by the omentum only after peritoneal inflammation was triggered by Zymosan-A, and this could not be recapitulated *ex vivo* (Fig. 2E). This suggested that CXCL1 was constitutively produced by the omentum, while the initial secretion of CCL2 and CXCL10 was dependent on the early recruitment of immune cells upon sensing of an inflammatory signal. The importance of radiation resistant stromal derived CXCL1 in the control of bacterial infection during peritonitis was previously demonstrated (Jin et al., 2017). Wholemount immuno-fluorescence staining of the omentum revealed that the expression of CXCL1 was particularly high in FALC cover cells (Fig. 2F). Given the spatially constrained expression of CXCL1 over the surface of FALCs and that neutrophils use FALC HEVs to enter the peritoneal cavity (Buscher et al., 2016), we next assessed whether CXCL1 was important for the recruitment of neutrophils to omFALCs during peritonitis.

### CXCL1 mediates active recruitment of neutrophils into omFALCs

To characterise the dynamics of neutrophil recruitment to omFALCs during peritonitis we used the well-established murine model, Zymosan-A induced peritonitis. Analysis of the single cell suspension of the omentum showed that peritonitis led to a transient increase in the number of Ly6G^+^ neutrophils in the omentum, peaking at 24 hours (Fig. 3A, Suppl. Fig. 4). Wholemount immunofluorescence staining and confocal analysis revealed that FALCs were the site of extensive recruitment of Ly6G^+^ Myeloperoxidase^+^ (MPO) neutrophils, which formed dense cellular aggregates between 6 and 24 hours post-induction of peritonitis (Fig. 3B,C). Estimation of the mean volume of FALCs during the course of peritonitis showed that the size of omFALCs increased exponentially during the first 18 hours post-Zymosan injection (75 fold), before rapidly contracting back to their normal size by 24 hours (Fig. 3D). This coincided exactly with the influx of MPO^+^ neutrophils and their disappearance by 24 hours (Fig. 3C,D). This accretion of neutrophils was specific to omFALCs and did not happen on the rest of the surface of the omentum, the parietal wall (Suppl. Fig 5) or the visceral mesothelium. We then tested if CXCL1 was involved in the accumulation of neutrophils at omFALCs during peritonitis, using an anti-CXCL1 blocking antibody. Eighteen hours post-Zymosan injection, CXCL1 blockade led to a 2.6-fold decrease in the number of neutrophils recovered from the omentum compared to mice treated with isotype control antibodies (Fig. 3E). Importantly, there was no difference in the number of neutrophils recovered from the peritoneal cavity, suggesting that the trafficking of neutrophils through omFALC HEVs was not altered (Fig. 3E). We thus concluded that the extensive recruitment of neutrophils we observed in FALCs was not just a consequence of increased trafficking of neutrophils through HEV but the result of active retention dependent upon CXCL1.

**Figure 3:**
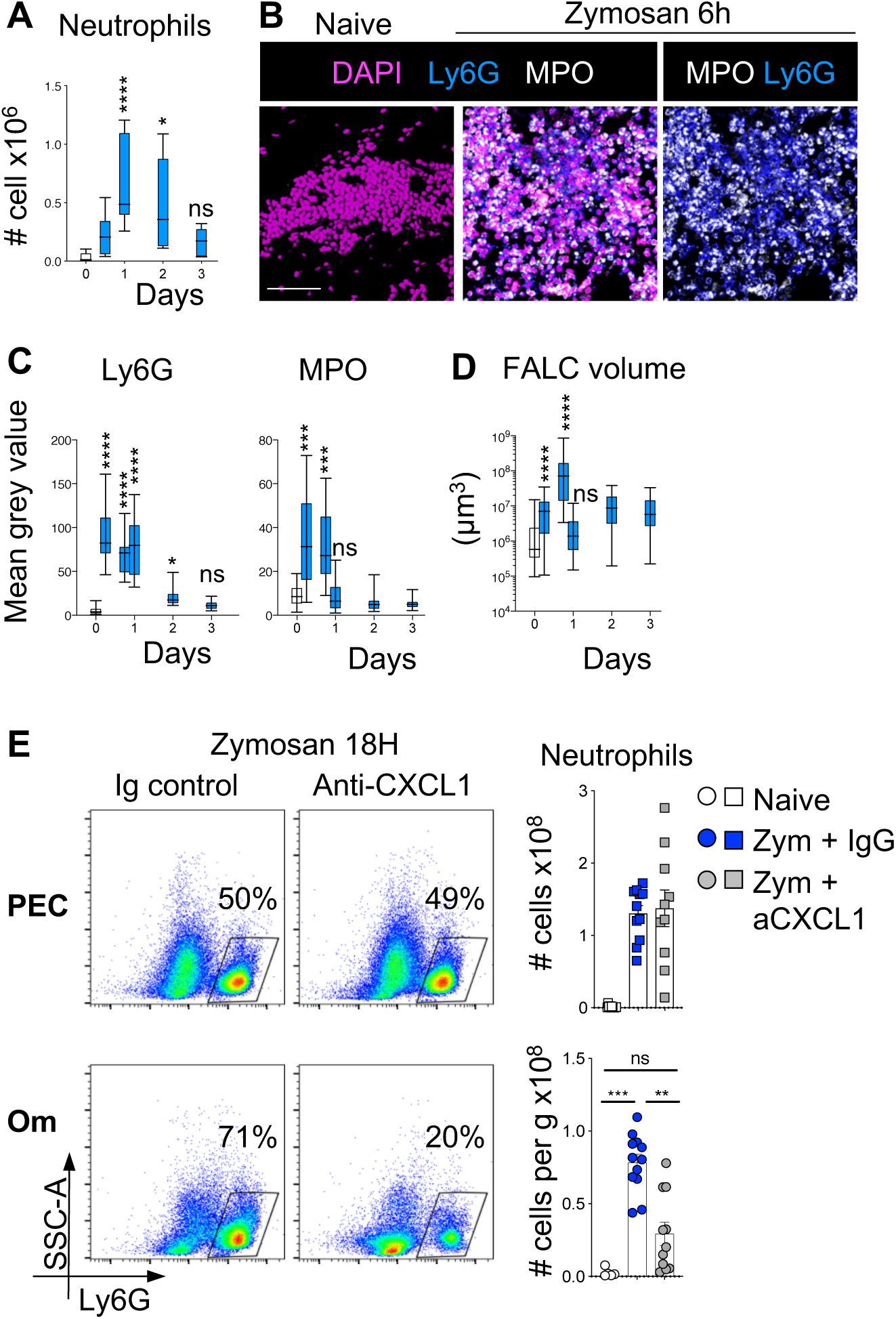
CXCL1 is required for the recruitment of neutrophils into omFALCs. **A** Number of neutrophils in digested omentum as assessed by flow-cytometric analysis (Supplementary Fig. 4) of naïve (white) and at the indicated time points following i.p. injection of 0.5mg Zymosan-A (blue). **B**, Representative confocal images of wholemount immuno-fluorescence staining of omentum from naïve and 6 hours post-Zymosan mice, DAPI (magenta), MPO (White) and Ly6G (Blue). Scale bar 50 μm. **C,** Quantification of the mean grey value of Ly6G and MPO stained as in **B** of omenta from naïve (white) and at the indicated time points following i.p. injection of Zymosan (blue). **D,** Quantification of the volume of omFALCs as assessed by measuring the maximal length, width and depth of clusters visualized with DAPI at the indicated time points following i.p. injection of 0.5mg Zymosan-A (blue). **E**, Representative density plots showing the proportion of neutrophils found in the PEC and omentum 18h after Zymosan i.p injection, in combination with the injection of either anti-CXCL1 or isotype control antibodies after 2h and number of neutrophils found in PEC and per g of omentum in naïve and treated mice. Data pooled from two independent experiments n=5-11 mice per group. Error bars show SEM. Box and whiskers showing min to max value. Kruksal Wallis test with Dunn’s multiple comparisons test or ANOVA with Sidak’s multiple comparisons test were applied after assessing normality using D’Agostino & Pearson Normality tests, ns= non-significant, **=P <0.01, *** P=<0.001, **** P=<0.0001.

### Neutrophils form large aggregates in omFALCs which are encapsulated in NET-like structures

Immunofluorescence imaging analysis of accumulated neutrophils within omFALCs revealed the presence of multiple areas of DAPI staining presenting with a non-nuclear, non-globular appearance (Fig. 4A, right panel). Neutrophils have the capacity to form neutrophil extracellular traps (NETs), which consist of uncoiled DNA scaffold decorated with proteases and anti-microbial peptides found in neutrophil granules. NETs are formed as a defence mechanism to immobilize invading microorganisms but also in response to sterile triggers (Boeltz et al., 2019; Papayannopoulos, 2018). In some conditions, NETs have been shown to mediate neutrophil aggregation (Leppkes et al., 2016; Munoz et al., 2019; Schauer et al., 2014). Chromatin decondensation, which is required for the formation of NETs can be mediated by the citrullination of histones. Staining for citrullinated histone H3 (CitH3), revealed that the areas of DAPI staining presenting with an extruded DNA pattern were highly stained for CitH3 (Fig. 4A). CitH3^+^ DNA was found in FALCs between 6 and 18 hours post-Zymosan injection at the peak of neutrophil recruitment (Fig. 4B and Fig 3A,C). The highest density of CitH3^+^ DNA was found on the surface of the expanded FALCs, which typically formed a dense core of neutrophils coated with a CitH3^+^ DNA outer-layer (Fig. 4C and Suppl. Fig. 6).

**Figure 4.**
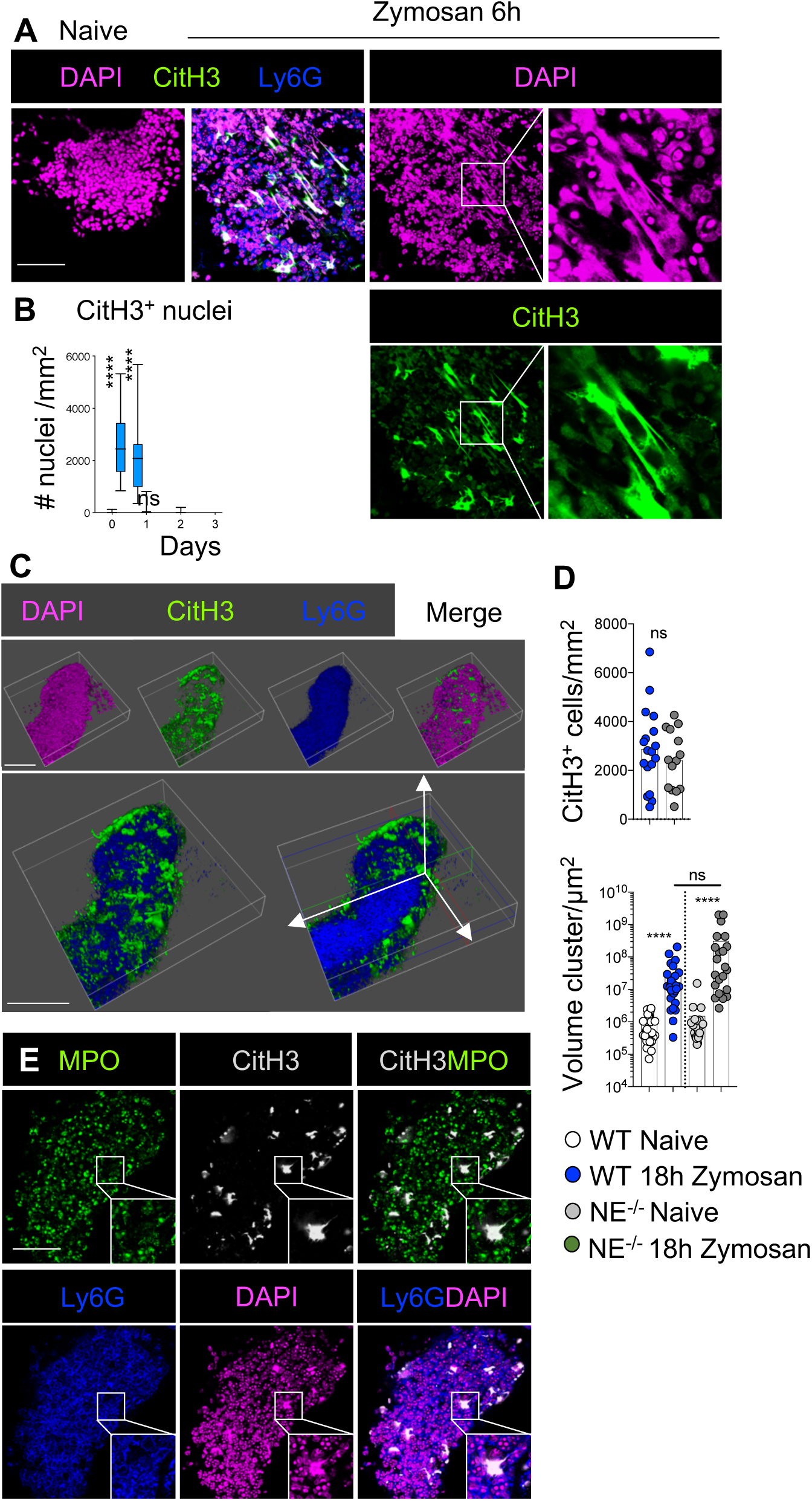
CitH3+ DNA coats neutrophil aggregates on omFALCs during peritonitis. **A**, Representative confocal images of wholemount immuno-fluorescence staining of omentum from naïve and 6 hours post-Zymosan mice, DAPI (magenta), CitH3 (Green) and Ly6G (Blue). Scale bar 50 μm. **B,** Quantification of the number of CitH3 positive nuclei per mm at the times indicated post Zymosan injection. Data pooled from two independent experiments n=5-11 mice per group. **C,** 3D reconstruction of omFALC obtained by confocal imaging of wholemount immunofluorescence staining of the omentum at 6 hours post-Zymosan injection showing the surface of a cluster (first row) and a clipped view of the inside of the cluster (second row). Ly6G (Blue), CitH3 (Green), DAPI (Magenta). Scale Bar 100μM. **D.** WT or NE^−/−^ mice were left naïve (WT in white, NE^−/−^ in grey) or injected i.p. with Zymosan (WT in blue, NE^−/−^ in grey) and omenta were collected 18 hours post-injection; the number of CitH3^+^ cells per mm and volume of each cluster was assessed by wholemount immunofluorescence staining and confocal analysis of omenta from naïve and treated mice. Data pooled from two independent experiments n=4 mice per group **E**. Representative confocal images of wholemount immuno-fluorescence staining of omentum isolated at 6H following Zymosan injection showing MPO (Green), Ly6G (Blue), CitH3 (White) and DAPI (Magenta), Scale Bar = 50μM. Student’s T-test, Kruksal Wallis test with Dunn’s multiple comparisons test or Mann Whitney test were applied after assessing normality using D’Agostino & Pearson Normality test, ns= non-significant, **** P=<0.0001.

To further characterise the CitH3^+^ DNA outer-layer and determine if it could be considered as NETs, we analysed the requirement for Neutrophil elastase (NE). NE is a granule serine protease which translocate to the nucleus, where it promotes chromatin decondensation and the formation of NETs (Papayannopoulos et al., 2010). Here we found that the formation of the CitH3^+^ DNA layer coating omFALC neutrophil aggregates and the expansion of these structures during Zymosan induced peritonitis was not affected in NE^−/−^ mice compared to WT mice (Fig. 4D). MPO, another granule serine protease, synergises the action of NE and is normally found associated with NETs (Papayannopoulos et al., 2010). Staining for MPO revealed that the CitH3^+^ DNA covering the neutrophil aggregates was not associated with MPO suggesting that MPO relocation to the nucleus was not required for the formation of the CitH3^+^ DNA outer-layer (Fig. 4E). Taken together, our data indicated that the formation of the CitH3^+^ DNA layer coating the omFALC neutrophil aggregates that form during Zymosan-induced peritonitis were independent of NE and thus different from the formation of “classical” NETs.

### Inhibition of PAD4 prevents the aggregation of neutrophils in omFALCs and the trapping of Zymosan by omFALCs

The enzyme PAD4, which mediates the conversion of arginine into citrulline, has been implicated *in vitro* and *in vivo* in the formation of NETs. The role of PAD4 in NET formation remains controversial and seems to be context dependent (Boeltz et al., 2019; Konig and Andrade, 2016). Recent work implicated PAD4 in the release of NETs and the capture of ovarian cancer metastasis by the omentum (Lee et al., 2019). In order to test whether PAD4 was involved in the capture of particulate contaminants and the formation of neutrophil aggregates on omFALCs we used fluorescently labelled Zymosan (Fluo-Zym) and the specific PAD4 inhibitor GSK484 (Lewis et al., 2015). Observation of the omentum with a stereo-microscope at 6 hours post-injection, revealed that omFALCs underwent a massive expansion and very effectively captured and concentrated Fluo-Zym particles. PAD4 inhibition completely abrogated both the omFALC expansion and the capture of Fluo-Zym particles by omFALCs (Fig. 5A-C). Confocal imaging showed that Fluo-Zym injection led to the formation of large neutrophil aggregates embedded in CitH3^+^ DNA, while PAD4 inhibition blocked the recruitment of Ly6G^+^ neutrophils into omFALCs, and significantly reduced the CitH3 staining compared to Fluo-Zym only controls (Fig. 5B,D). We confirmed that PAD4 inhibition was associated with omFALCs failing to capture Fluo-Zym particles and expand in size (Fig. 5A,E). In contrast, we found that GSK484 did not block the entrance of neutrophils into the peritoneal cavity since we recovered twice as many neutrophils from the peritoneal cavity when mice received GSK484 and Fluo-Zym compared to Fluo-Zym only (Fig. 5F). Taken together, these results indicated that GSK484 inhibited neutrophil accretion in omFALCs, which led to a very severe reduction in the capacity of the omentum to capture Zymosan particles (Fig. 5A). While neutrophils clearly underwent citrullination of Histone H3 during Zymosan induced peritonitis, the fact that PAD4 inhibition led to a near complete abrogation of neutrophil aggregation within FALCs did not allow us to conclude on whether PAD4 was involved in the formation of the CitH3^+^ DNA outer-layer we observed.

**Figure 5:**
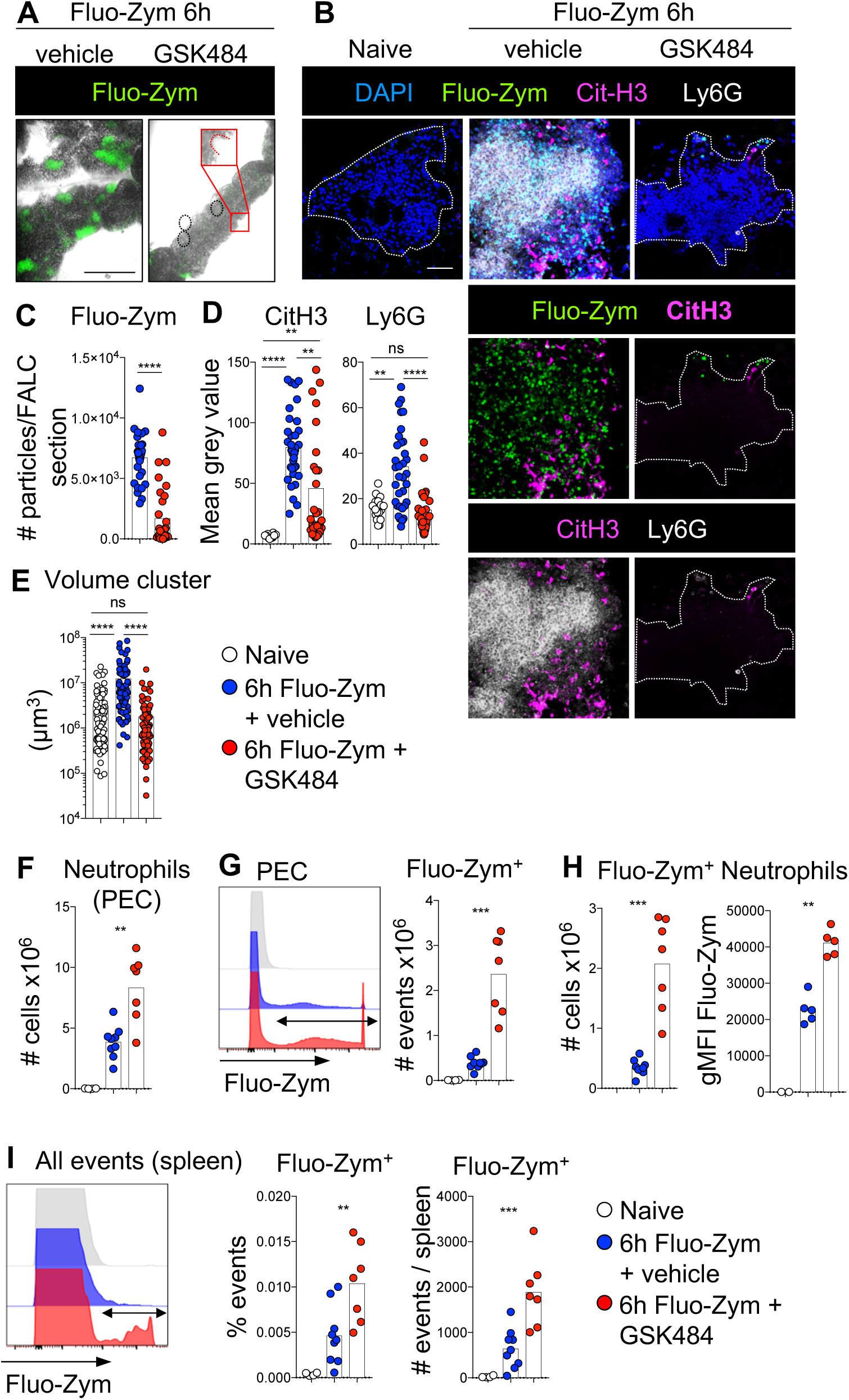
Inhibition of PAD4 prevents neutrophil aggregation and the capture of Zymosan particles within omFALCs, while increasing the retention of Zymosan in the peritoneal cavity and its spread to the spleen. **A-I,** Mice were left naïve (white circles) or injected i.p. with Fluo-Zym in combination with the PAD4 inhibitor GSK484 (red circles) or vehicle (blue circles) and the omentum, PEC and spleen were analyzed 6 hours post injection. Representative low (**A,** scale bar 1 mm) and high (**B,** scale bar 50 μm) magnification confocal images of wholemount immuno-fluorescence staining of omentum. Number of Fluo-Zym particles per mm^2^ of omFALCs (**C**), mean grey value for CitH3 and Ly6G staining (**D**), volume of omFALCs as assessed by measuring the maximal length, width and depth of clusters visualized with DAPI (**E**). Total number of PEC neutrophils (**F**). Representative histogram plot showing fluorescence intensity of Fluo-Zym in all PEC and number of Fluo-Zym^+^ PEC (**G**). Number of Fluo-Zym^+^ neutrophils and MFI of Fluo-Zym within Fluo-Zym^+^ neutrophils (**H**). Representative histogram plot showing fluorescence intensity of Fluo-Zym in spleen and percentage and number of Fluo-Zym^+^ events per spleen (**I**). Data pooled from two independent experiments with n=7-8 mice per group. Kruksal Wallis test with Dunn’s multiple comparisons test or Mann Whitney test were applied after assessing normality using D’Agostino & Pearson Normality test, ns= non-significant, **=P <0.01, *** P=<0.001, **** P=<0.0001.

### Inhibition of PAD4 leads to impaired clearance of Zymosan particles from the peritoneal cavity and increases dissemination to the spleen

We next addressed the effect of PAD4 inhibition on peritoneal neutrophil clearance of Zymosan-A and discovered that in addition to increasing peritoneal neutrophil retention (Fig. 5F), inhibition of PAD4 resulted in significantly increased retention of Fluo-Zym particles within the peritoneal cavity (Fig. 5G). There was a three-fold increase in the proportion of neutrophils retaining Fluo-Zym particles, with a near two-fold increase in Fluo-Zym MFI, indicating that PAD4 inhibition led to increased phagocytosis of Fluo-Zym particles by peritoneal neutrophils (Fig. 5H). Finally, we found that PAD4 inhibition led to increased dissemination of Fluo-Zym particles to the spleen (Fig. 5I). Collectively, these results indicated that PAD4 dependent neutrophil aggregation within omFALCs provided a rapid and efficient mechanism to clear particulate material from the peritoneal cavity and limit the systemic spread of peritoneal contaminants.

### Neutrophil recruitment and aggregation confer increased adhesive properties to omFALCs

In the peritoneal cavity, the capture of particles by omFALCs results from a combination of peritoneal fluid flow through the omentum and the capacity of the omentum to retain these particles. To assess the importance of neutrophil aggregation on the adhesive properties of the omentum independent of fluid flow we measured *ex vivo* the capacity of the omentum to capture bacteria relevant to peritonitis by briefly incubating fluorescently-labelled *E.coli* bioparticles with omentum samples isolated from untreated control mice, or from mice undergoing Zymosan induced peritonitis that had been treated with isotype control antibodies or anti-CXCL1 antibodies *in vivo* (Fig. 6A). Omenta were collected at 18 hours post-Zymosan injection and incubated *ex vivo* for 5 minutes at room temperature with *E. coli bioparticles*, before fixation and wholemount immuno-fluorescence imaging. Omentum tissue sampled from mice with peritonitis proficiently trapped *E.coli* whereas those from control mice without peritonitis did not (Fig. 6A). Blockade of CXCL1 resulted in fewer neutrophils within the omentum and a failure to efficiently trap *E. coli,* indicating that CXCL1-mediated recruitment of neutrophils to the omentum dramatically increased the capacity of the omentum to capture bacterial contaminants. We confirmed the importance of neutrophil recruitment and aggregation for the increased adhesive properties of the omentum during peritonitis using omenta isolated from naïve mice, those undergoing peritonitis and those undergoing peritonitis with PAD4 inhibition, and *E.coli* expressing the fluorescent protein mCherry. Omenta from naïve mice did not trap *E.coli*, contrary to omenta from mice with peritonitis (containing NET-like structures) where large areas of FALCs were covered in *E.coli* (Fig. 6B). PAD4 inhibition led to a marked decrease in the efficiency with which the omentum captured *E. coli* (Fig. 6B). These data suggest that neutrophil aggregation within omFALCs during peritonitis contributes to omental clearance of peritoneal bacterial contaminants.

**Figure 6.**
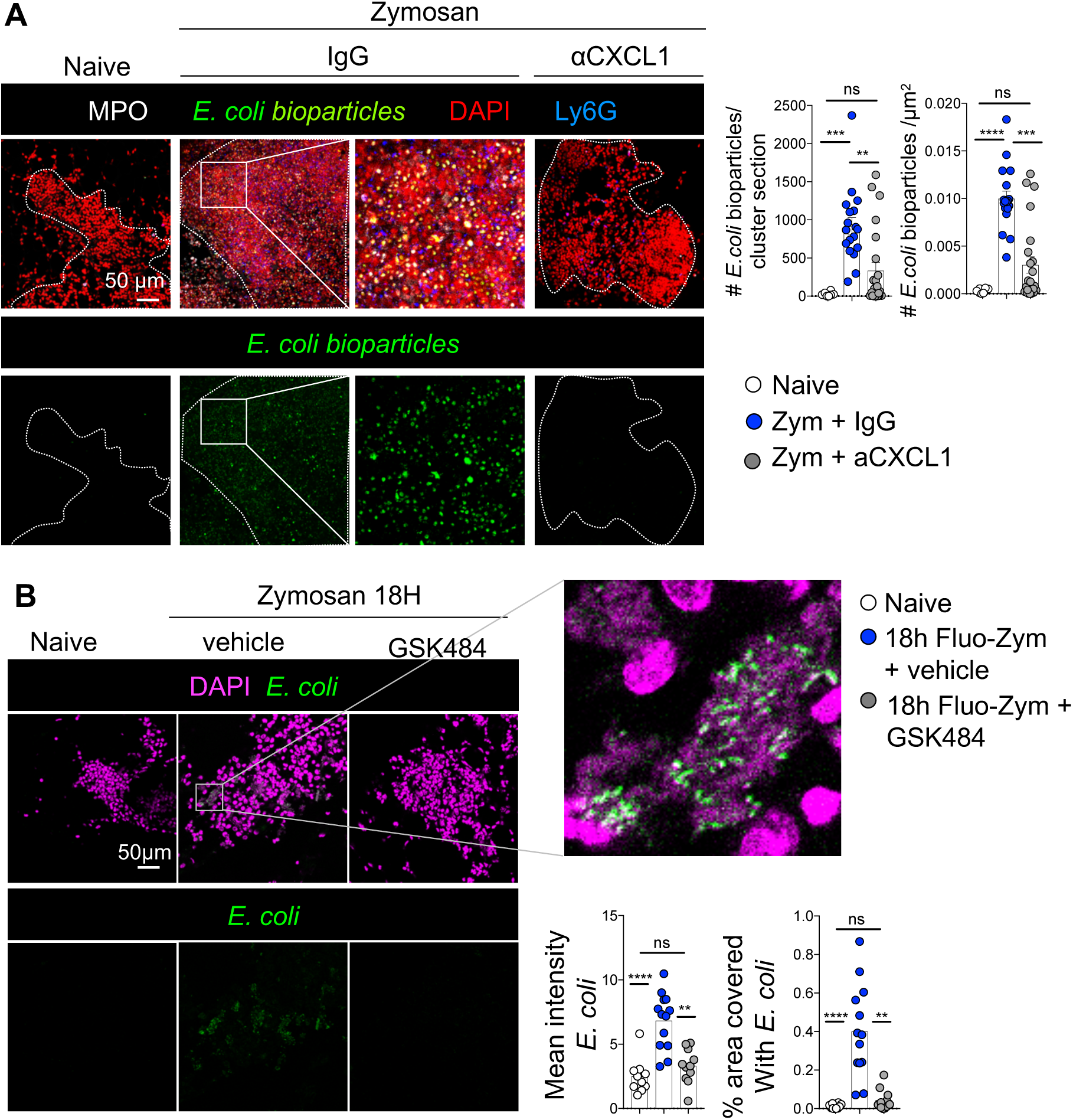
CXCL1 and PAD4 are required for the adhesion of bacteria to the omentum in the absence of fluid flow. **A**. Representative confocal images of wholemount immuno-fluorescence staining of Omenta isolated from either naïve animals or 18h after Zymosan i.p injection, in combination with the injection of either anti-CXCL1 or isotype control antibodies after 2h were cultured with fluorescently labelled *E.coli* bio-particles for 10 minutes, number of *E.coli* bioparticles found per cluster section and per μm^2^ of cluster were graphed. Scale bar 50 μm. MPO (White) *E.coli* (Green), DAPI (Red), Ly6G (Blue). Data pooled from two independent experiments n=10-11 mice per group. Error bars show SEM. Kruksal Wallis test with Dunn’s multiple comparisons test or ANOVA with Sidak’s multiple comparisons test were applied after assessing normality using D’Agostino & Pearson Normality tests, ns= non-significant, **=P <0.01, *** P=<0.001, **** P=<0.0001.**B**, Mice were left naïve (white circles) or injected i.p. with Zymosan in combination with the PAD4 inhibitor GSK484 (grey circles) or vehicle (blue circles), the omenta were collected 18 hours post-injection and incubated *in vitro* for 5 minutes with mCherry^+^ *E. coli*. Representative confocal images of wholemount immuno-fluorescence staining of omentum. Scale bar 50 μm. Mean grey value of *E. coli* in omFALCs and percentage area covered by *E. coli.* Data pooled from two independent experiments with n=7-8 mice per group. Kruksal Wallis test with Dunn’s multiple comparisons test or Mann Whitney test were applied after assessing normality using D’Agostino & Pearson Normality test, ns= non-significant, **=P <0.01, *** P=<0.001, **** P=<0.0001. Scale bar 50 μm.

### Neutrophils are recruited to the human omentum during peritonitis

We reasoned that appendicitis would provide a useful translational platform to examine the function of the omentum during peritonitis. During acute appendicitis the omentum wraps itself around and adheres to the inflamed appendix (Morison, 1906). By comparison, in patients undergoing laparoscopic surgery for biliary colic (surgery for gallstones without active inflammation) the omentum and appendix are both uninvolved. We recruited patients who were undergoing laparoscopic surgery for possible or suspected acute appendicitis (App), or laparoscopic cholecystectomy for biliary colic (non-inflamed control group; NI) (Suppl. Table 1). We found that acute appendicitis induced a stark influx of CD19^−^CD3^−^NCAM^−^CD15^+^ neutrophils into the omentum and the peritoneal cavity (Fig. 7A,B).

**Figure 7:**
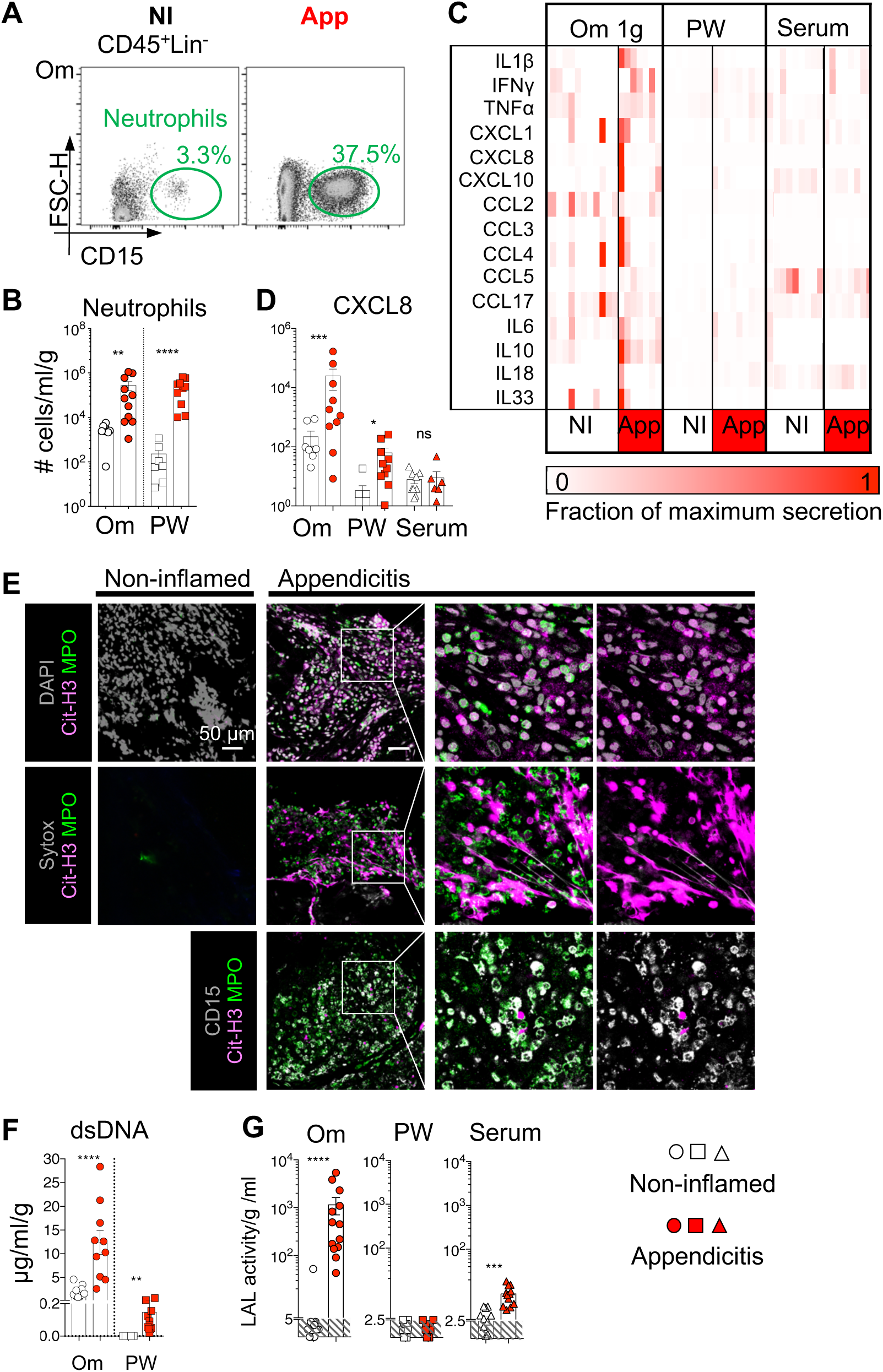
The human omentum recruits neutrophils and collects bacterial antigens during appendicitis. **A**, Representative gating strategy showing CD45^+^ CD19^−^, CD3^−^, NCAM^−^, CD15^+^ neutrophils in the omentum (Om) of non-inflamed (NI) and appendicitis (App) patients. **B,** Number of neutrophils found per gram of Om or per ml of PW of NI and App patients. Patients were stratified based on surgical outcome into one of 2 groups, NI (white) or App (red), n=7 and 10 patients per group. Error bars show SEM. **C,** Inflammatory chemokines & cytokines found in 2h Om explant culture supernatant, PW and serum comparing NI and App patients. **D,** Amounts of CXCL8 per g per ml of 2h Om explant culture supernatant **E**, Representative confocal images of wholemount immunofluorescence staining of omentum biopsies from non-inflamed control or appendicitis patients (n≥6) showing in grey DNA stained with DAPI (upper) or extra-cellular DNA stained with SYTOX (middle, grey), or CD15 (lower); in magenta CD11b; in green Myeloperoxidase (MPO) and in red Citrullinated histone H3 (CitH3). **F**, Amounts of double stranded DNA (dsDNA) released into the supernatant of 2h omental explant culture per g and per ml and per ml of peritoneal wash (PW) in control and appendicitis patients. **G**, LAL activity within the supernatant of 2h omental explant culture per g and per ml and per ml of PW and serum in control and appendicitis patients. n=10-13 patients per group. Error bars show SEM. Unpaired student’s T-test or Mann Whitney test were applied after assessing normality using D’Agostino & Pearson and Shapiro-Wilk Normality tests n.s.=non-significant, * = P<0.05, **=P <0.01, *** P=<0.001.

We next investigated whether the mechanisms regulating recruitment of neutrophils to the human omentum were similar to mouse by analysing the secretion of inflammatory chemokines and cytokines over a two-hour interval in *ex vivo* human omentum explant cultures (Fig. 7C). CXCL8 (the human homologue to murine CXCL1) and IL-1β, two important factors for neutrophil recruitment and activation (Biondo et al., 2014), were secreted at much higher levels in omental explant cultures and peritoneal lavage in inflammatory conditions compared to controls, but not in serum (Fig. 7C, D). Strikingly, we found that in patients with acute appendicitis, 1g of omental explant released 25ng of CXCL8, approximately 400 times more than the amount found per ml of peritoneal lavage, despite the number of neutrophils being similar in equivalent volumes of omentum and wash fluid (Fig. 7D) indicating that the human omentum is a key site of CXCL8 release and neutrophil recruitment during peritonitis.

### CitH3^+^ NET-like structures are released on the human omentum during peritonitis

Finally, we tested whether neutrophils recruited to the omentum during peritonitis in humans also released CitH3^+^ NET-like structures. We performed wholemount immunofluorescence staining and detected the presence of extracellular DNA fibrils using SYTOX or DAPI staining, which we found to co-localise with CitH3, in regions where there was a substantial CD15^+^ neutrophil infiltrate during acute appendicitis (Fig. 7E). We could also detect areas of CitH3^+^ DNA staining which did not co-localise with MPO which was comparable to the murine omentum during peritonitis. No NET-like structures were detected in any non-inflamed control samples (Fig. 7E). In addition, we detected dsDNA in omental explant culture supernatants and in the peritoneal wash fluid from patients with acute appendicitis, but not from control patients (Fig. 7F). Since an equivalent number of neutrophils were present per gram of omentum and per ml of peritoneal wash (Fig. 7B), the fact that dsDNA was released in the greatest amounts by the omental explants strongly suggests that the omentum supports DNA release from neutrophils. Using a limulus amoebocyte lysate (LAL) assay, we measured bacterial LPS within omental explant cultures and detected LPS release into culture supernatants from omentum isolated from patients with acute appendicitis, but not from non-inflamed patients. Patients with appendicitis also had higher serum levels of LPS, but there was no evidence of LPS within the peritoneal wash (Fig. 7G). This suggested that during appendicitis, neutrophil aggregation and the release of CitH3^+^ NET-like structures on the omentum mediated successful capture of bacteria/bacterial antigens arising from the inflamed appendix and thus protected the wider peritoneal cavity from contamination and generalized peritonitis.

## Discussion

In the present study, we uncover new facets of the stromal-immune cell interactions which govern omFALC function within the peritoneal cavity and reveal how neutrophils mediate the clearance of peritoneal contaminants by omFALCs. We show that the surface of FALCs is covered by differentiated mesothelial cells specialising in the secretion of inflammatory mediators (*Cxcl13^+^* FALC cover cells) and the response to virus (*Ifit^+^* FALC cover cells). We demonstrate that during peritonitis, neutrophils rapidly accumulate within omFALCs where they form large aggregates concentrating peritoneal contaminants and limiting their systemic translocation. We establish that the formation of these aggregates depends on two mechanisms: (i) CXCL1, which is produced by Cxcl13^+^ FALC cover cells; and (ii) the PAD4 dependent formation of a CitH3^+^ DNA outer-layer coating the omFALC neutrophil aggregates (Supplementary Fig. 6). In humans with appendicitis, we validate that the omentum is also a site of neutrophil recruitment and the release of CitH3^+^ DNA, and we provide evidence that the omentum efficiently captures bacterial antigens, leaving the peritoneal cavity free of contamination.

The mesothelium is a critical component of adipose tissue, source of mesenchymal and adipocyte progenitors and acts as a physical barrier to the serosal cavity. In the omentum the mesothelium exerts an additional function: the filtration of peritoneal fluid which is facilitated by the presence of stomata (Mironov et al., 1979) that afford particles and cells entrance into FALCs and enable fluid to be drained into neighbouring lymphatic vessels. Our analysis revealed the exquisite adaptation of the mesothelium for this function, which gives rise to *Cxcl13^+^* FALC cover cells, specialised in the attraction of immune cells and the secretion of inflammatory mediators and *Ifit^+^* FALC cover cells competent for the secretion of anti-viral factors.

Until recently, due consideration hasn’t been paid to the issue of how immune cells perform their function in the liquid phase of the serous cavities. Recent work revealed that resident peritoneal macrophages undergo a clotting response within the first 2h of contamination of the peritoneal cavity, providing a first mean of clearing peritoneal contaminants via coagulation and adhesion (Zhang et al., 2019). This mechanism serves to convert fluid phase inflammation to a solid state within the clots. Neutrophil aggregation within FALCs also enable the conversion to a solid state which is required for efficient clearing of particles from the fluid phase. In doing so, neutrophils provide a timely relay system for the neutralisation of peritoneal contaminants, since resident peritoneal macrophages are rapidly sequestered in clots (Zhang et al., 2019). In gout and pancreatitis, presence of high density neutrophils in combination with the release of NETs lead to the formation of large DNA aggregates. In gout, these aggregated NETs have the ability to degrade cytokines and chemokines via serine proteases and may be important to limit inflammation(Schauer et al., 2014). In pancreatitis, PAD4 mediates the release of NETs which cause neutrophil aggregation & occlusion of pancreatic ducts (Leppkes et al., 2016). Most recently, aggregated NETs have been implicated in the formation of gallstones (Munoz et al., 2019); Munoz *et al* report that macropinocytosis of small crystals results in NET release that is dependent upon both PAD4 and ROS. ROS dependent formation of aggNETs has also been reported to provide an anti-inflammatory shield around necrotic neutrophil filled cores in the context of nanodiamond induced sterile inflammation (Biermann et al., 2016). Here we show that omFALCs are also sites where high densities of actively phagocytosing neutrophils aggregate in a PAD4 dependent mechanism during peritonitis.

Previous studies have posited that both MPO (Metzler et al., 2011) and NE are necessary for NET release and must translocate into the nucleus in order to mediate the decondensation of chromatin and the release of bonafide NETs (Papayannopoulos et al., 2010). In this work we find *in vivo* during peritonitis that neutrophils aggregate within, and CitH3^+^ DNA accumulates over the surface of expanded omFALCs even in the absence of NE suggesting a novel mechanism of DNA release that does not conform to the same rules as classical NETosis. In omFALCs during peritonitis, CitH3^+^ DNA did not robustly co-localise with MPO staining, again indicating a non-classical mechanism of neutrophil CitH3+ DNA release. Furthermore, we found CitH3^+^ DNA extrusions on the surface of the human omentum during appendicitis, re-confirming the presents of NETs in the human peritoneal cavity as shown by others in similar & diverse disease contexts (Brinkmann et al., 2004; Lee et al., 2019). In contrast to NE, we found via delivery of a chemical inhibitor GSK484 *in vivo* that the formation of omFALC neutrophil aggregates is dependent upon PAD4. This work contributes to mounting evidence that mechanisms of NET release vary dependent upon context and location (Boeltz et al., 2019); an unsurprising finding given the exquisite complexities of the immune response.

We hypothesise based on our results (Fig. 8) that following the initial FALC Cover cell CXCL1 mediated recruitment of zymosan-loaded peritoneal neutrophils to omFALCs, the release of extracellular DNA facilitates the adhesion of further waves of neutrophils which also extrude their DNA resulting in the large aggregates imaged at 6-18h. To account for the staining pattern seen, we assume that re-modelling of the DNA extrusions must occur to clear CitH3 from the centre of the aggregates while leaving the fluid-facing aggregates coated in NET-like structures. This intriguing CitH3^+^ DNA outer-coating raises multiple questions, 1) What is the role of the CitH3^+^ DNA on the fluid facing surface of the cluster; are CitH3^+^ neutrophil aggregated omFALCs anti-bacterial? 2) Are the aggregations performing a purely structural role by barricading the omFALC to limit the spread of peritoneal contaminants? 3) How are the aggregates cleared during the resolution of peritonitis? 4) Does resolution require exposure of CitH3^+^ DNA?

**Figure 8:**
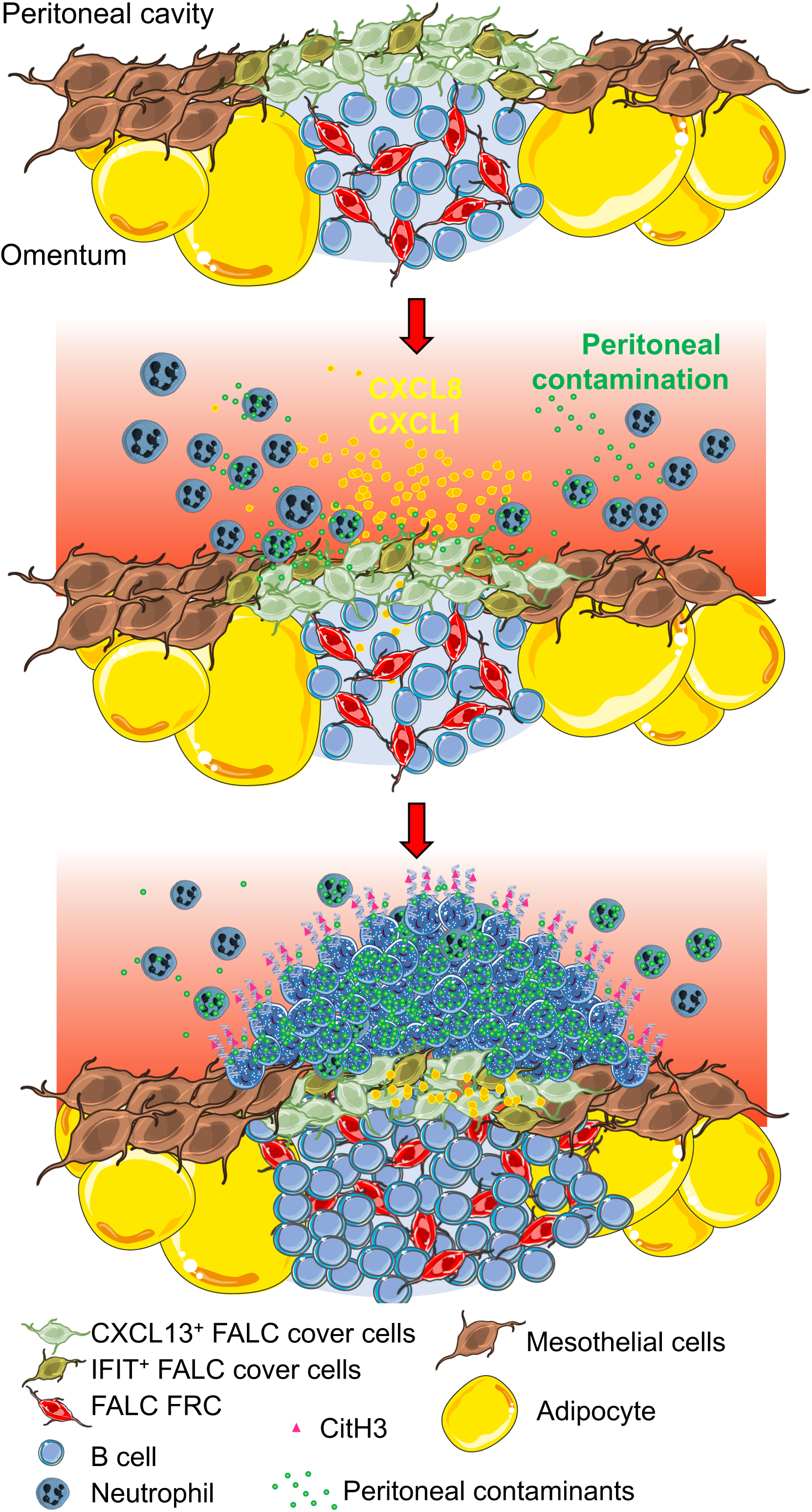
Model illustrating the mechanisms underlying the recruitment of neutrophils to the omentum during peritonitis. FALC stroma is made up of a network of FALC FRCs interacting with B cells and of two distinct subset of FALC cover cells, one specialised in the secretion of inflammatory mediators (CXL13^+^ FALC cover cells) and one specialised in the response to virus (IFIT^+^ FALC cover cells). Upon sensing of danger or pathogen, CXCL13^+^ FALC cover cells release CXCL1, triggering an intense recruitment of neutrophils, the release of CitH3^+^ DNA and neutrophil aggregation. The formation of neutrophil aggregates covered in CitH3^+^ DNA is required for the capture of particulate contaminants by omFALCs and increases its adhesivity to bacteria protecting the wider peritoneal cavity from contamination and generalized peritonitis.

In sepsis, NETs have been shown to promote the toxic inflammatory and procoagulant host response to endotoxin (Martinod et al., 2015). However, our data points toward an important role for omental NET release to limit the propagation of contaminants from the peritoneal cavity to the circulation. Due to constant drainage of peritoneal fluid through FALCs and their high vascularisation (Buscher et al., 2016; Dickinson, 1906; Gray et al., 2012), FALCs are the main portals for the transit of peritoneal contaminants systemically. While targeted NET release on omFALCs is beneficial during peritonitis, it may play a detrimental role in ovarian cancer metastasis (Lee et al., 2019). Targeting FALC cover cells may prove a useful approach to limit omental metastasis.

## Acknowledgments

We are extremely grateful to: The Edinburgh Emergency Surgery Study Group, Royal Infirmary of Edinburgh, NHS Lothian: Graeme Couper, Christopher Deans, Gavin G P Browning, Anna M Paisley, Bruce Tulloh, Richard J E Skipworth, Simon Paterson-Brown, Rajan Ravindran, Andrew de Beaux, Stuart McKechnie, Andrew Healey, Dimitri Damaskos, Marina Garcia and all Royal Infirmary surgical trainees for facilitating intraoperative sample collection. The Wellcome Trust Clinical Research Facility, NHS Lothian, especially Beena Poulose, for clinical study support. The CALM facility for advice on flow cytometry and confocal microscopy. Stephen Mitchell for assistance with SEM. Central Bioresearch Services, especially Gidona Goodman, Chris Flockhart and Will Mungall for their technical support. D. Leach (University of Edinburgh) for the RecA-mCherry *E.coli.* Figure 8 was compiled using images from Servier Medical Art.

## Author contributions

**LJJ** Conceptualization, Investigation, Project administration, Formal analysis, Writing-original draft, Visualization. **PS, MMP, KM, JP**, **NH,** Investigation, Analysis and review. **BH, RD, MN** Investigation. **DJM** Clinical protocol draft, obtaining ethical and clinical governance permissions, Chief Investigator (clinical arm), methodology, project administration, writing review/editing. **CB** Funding Acquisition, Supervision, Conceptualization, Investigation, Project administration, Formal analysis, Writing-original draft, visualization.

## Methods

### Animals and peritonitis models

All experiments were conducted under a license granted by the Home Office (UK) that was approved by the University of Edinburgh animal welfare and ethics review board. All individual experimental protocols were approved by a named veterinarian surgeon prior to the start of the experiment. Experiments were performed using female C57BL/6 (C57BL/6JOlaHsd) or NE deficient mice (Belaaouaj et al., 1998) aged 8-12 weeks. All animals were bred and maintained under specific pathogen–free conditions at the University of Edinburgh Animal Facilities. For the peritonitis models, mice were injected intra-peritoneally with either 0.5mg Zymosan-A (Sigma) in PBS, 0.25mg of Fluorescein labelled Zymosan-A (Sigma), or PBS alone and samples were isolated 2-72h later. To block NETosis, mice were injected i.p. with 400μg/mouse of GSK484 (Cayman Chemical). Blocking antibodies against CXCL1 (Clone MA5-23745, Invitrogen) or isotype control IgG (eBioscience) antibodies were injected i.p. 2h after induction of peritonitis (40μg/mouse). Following Zymosan and anti-CXCL1 treatment, omenta were isolated and cultured with mCherry *E. coli* (Amarh et al., 2018) or *E. coli* bioparticles (Sigma) *in vitro* for 5 minutes prior to thorough washing with RPMI-1640 (Sigma), and wholemount staining as described below.

Peritoneal exudate cells (PEC) were isolated by flushing murine peritoneal cavities with RPMI 1640 (Sigma). Murine omenta were enzymatically digested with 1mg/ml Collagenase D (Roche) for 35 minutes at 37°C in RPMI 1640 (Sigma) containing 1% Fetal Bovine Serum (Sigma). Spleens were mechanically disrupted using glass slides.

### Human subjects

This study was approved by the Regional Research Ethics Committee (SE Scotland REC 02; 16/SS/0042), the University of Edinburgh/NHS Lothian ACCORD R&D Office (ref: 2016/0035) and the Office of the Caldicott Guardian, NHS Lothian (patient confidentiality advocate). Individuals were recruited after informed, signed consent was obtained. Clinical data and samples were collected from patients undergoing laparoscopic surgery under general anaesthesia for the following indications: biliary colic and possible or suspected appendicitis, at the Royal Infirmary of Edinburgh between 1^st^ April 2016 and 30^th^ June 2018. After the induction of anaesthesia, a single tube of blood was collected into a BD Vacutainer prefilled by the manufacturer with the anticoagulant EDTA. At operation, and as soon as practical and safe after the insertion of the laparoscopic ports (to avoid iatrogenic contamination with blood), 25mL of sterile 0.154 M NaCl solution was washed into the area of interest and then aspirated using a sputum trap interposed into the surgical suction equipment. Next a 5 cm^3^ sample of omentum was resected with scissors to avoid diathermy artefact, retrieved and the sample site haemostasis ensured with diathermy. A 2 cm^2^ sample of parietal peritoneum adjacent to a port site was obtained and haemostasis ensured. Operations then continued as planned. Clinical samples were handled as follows: omentum and parietal peritoneum samples were collected into 20 ml of sterile dPBS within a 50ml tube. Peritoneal washings were placed into a sterile 50ml tube. All patient samples were stored on wet ice or at 4 °C prior to collection from the clinical research facility and transported to the research laboratory on foot, samples collected after 5pm were processed the following day. If there were any unexpected findings at surgery, e.g. free peritoneal blood or peritoneal malignancy, patients were removed from the study and no tissue samples were taken for research purposes. There were no adverse effects due to the research study conduct.

### Human sample preparation

Human Omentum was weighed and 0.035 – 1.8g of tissue was digested using 2mg/ml Collagenase I (Worthington) in PBS (Invitrogen/sigma) 2% Bovine Serum Albumin (BSA, Sigma), samples were disrupted using an Octolyser (Miltenyi), incubated at 37°C with intermittent shaking for 45 minutes, subjected to a second Octolyser dissociation step, ions were chelated by addition of EDTA (0.5M, Sigma), samples were filtered through a 100μM filter (BD) and washed with 20ml of 2%BSA PBS prior to centrifugation at 1700pm for 10 minutes. The cell pellet was resuspended in 2ml of PBS 2% BSA for flow-cytometric analysis. Peritoneal wash and blood samples were centrifuged, the supernatant and serum were collected for further analysis and the cell pellet was resuspended for flow-cytometric analysis. Cell numbers and live cell count were determined using a BioRad TC20 automated cell counter and 0.4% Trypan Blue (Sigma). For the omentum *ex vivo* culture, a small piece of omentum (between 0.02 and 0.06g) was placed in culture in 1ml of RPMI 1640 (Sigma) 10% foetal bovine serum (Sigma) 2 mM L-glutamine (Sigma) for 2 hours at 37 °C.

### Flow cytometry

Murine cells were stained with LIVE/DEAD (Invitrogen), blocked with mouse serum and anti-murine CD16/32 (clone 2.4G2, Biolegend) and stained for cell surface markers (See Supplementary Table 2 for list of antibodies used). Human samples were blocked with serum, stained for cell surface markers (See Supplementary Table 2 for list of antibodies used), and DAPI was added to the cells prior to acquisition. All samples were acquired using a BD Fortessa and analyzed with FlowJo software (Tree Star).

### Detection of chemokines, cytokines, dsDNA and LPS

dsDNA was detected in lavage fluid and omentum culture supernatants using the picogreen assay following the manufacturer’s instructions (Invitrogen). LPS was detected in human serum, omentum culture supernatants and peritoneal lavage fluid using the ToxinSensor^TM^ Chromogenic LAL Endotoxin Assay Kit (GenScript) following manufacturer’s instructions. Human Pro-inflammatory chemokines, Human Inflammation and Murine CXCL1, CCL2 and CXCL10 Legendplex arrays (Biolegend) were used to detect cytokines and chemokines following the manufacturer’s instructions. For the heatmap representation in Fig. 6e, original values were scaled between 0 and the maximum value detected for each cytokine and presented as a fraction of maximum secretion.

### Wholemount immunofluorescence staining and confocal microscopy

Human and mouse omentum samples were fixed for one hour on ice in 10% NBF (Sigma) and then permeabilized in PBS 1% Triton-X 100 (Sigma) for 20 minutes at room temperature prior to staining with primary antibodies for one hour at room temperature in PBS 0.5% BSA 0.5% Triton. After washing in PBS, tissues were stained with secondary antibodies for one hour at room temperature in PBS 0.5% BSA 0.5% Triton. For extra-cellular DNA staining, human omental tissues were first stained with SYTOX Blue (Invitrogen) 1 in 5000 in RPMI 1640 (Sigma) for 30 minutes at room temperature, washed in RPMI and then fixed in 10% NBF (Sigma) prior to permeabilization and staining. Antibodies used are listed in Supplementary Table 2. After mounting with Fluoromount G, confocal images were acquired using a Leica SP5 or SP8 laser scanning confocal microscope. Image analysis was performed using Fiji and 3D reconstruction was created using LAS-X-3D (Leica). The mean grey value of MPO, Ly6G, CitH3 and mCherry *E. coli* was calculated inside a perimeter delimited manually as the border of the cluster. To calculate FALC volume, we manually assessed the maximum length, width and depth (z) of the clusters visualised with DAPI using 10x objective while scanning on SP5 confocal microscope. To calculate the area of FALCs covered by *E. coli,* the perimeter of the FALC was delimited manually and a fixed threshold for *E.coli* fluorescence was set. The number of CitH3^+^ nuclei and *E. coli* bioparticles was calculated using the “analyse particles” function of Fiji.

### Droplet based scRNAseq and data pre-processing

Immediately post-sorting, DAPI^−^CD45^−^Ter119^−^CD41^−^CD31^−^ stromal cell pooled from the omenta from three naive mice were run on the 10X chromium (10X Genomics) and then through library preparation following the recommended protocol for the Chromium Single Cell 3’ Reagent Kit (v2). Libraries were run on the NovaSeq S1for Illumina sequencing. Sequence reads were processed using the Cell Ranger v2.1.1 Single-Cell Software Suite from 10x Genomics. Reads were aligned to the mm10 mouse references genome. Genes were excluded if they were expressed in fewer than three cells. Cells were excluded on a number of criteria: those with fewer than 300 genes (123 cells), those with fewer than 300 or greater than 16000 UMIs (32 cells), those with mitochondrial gene proportion of over 20% of total UMI counts (18 cells) or those with a UMI:gene ratio of over 7:1 (4 cells). A global normalisation was performed where gene expression was normalised for each cell based on its total expression before being multiplied by a scale factor of 10,000 and log transformed. Variation in the UMI counts of each cell was regressed using a linear regression. Residuals from this model were centred and scaled by subtracting the average expression of each gene followed by dividing by the standard deviation of each gene. We obtained the transcriptional profile of 3,888 cells that passed quality control and filtering, for which an average of 2,303 genes per cells were measured.

### Dimensionality reduction, clustering, differential expression analysis and data visualisation

Dimensionality reduction, unsupervised clustering and differential gene expression were performed using *Seurat* R package version 2.3.4(Butler et al., 2018). Shared nearest neighbour (SNN) clustering was performed using principle components between 1 and 10, as determined by the dataset variability shown in the principle component analysis (PCA). The resolution parameter was optimised based on the number of resulting clusters. Clusters were initially categorised into cell lineages based on expression of known marker genes. Clusters annotated as endothelial (*Pecam1, Cdh5*), immune (*Ptprc*), proliferating (*Mki67, Pcna, Top2a*) or those with a median number of genes per cell below 1000 were excluded from further analysis. The final dataset contained 3888 cells and 15357 genes.

This cleaned dataset was re-clustered using the same methodology described above. Differential gene expression analysis was conducted using the *FindAllMarkers* function of *Seurat* and a Wilcox rank sum test (Supplementary Table 3). Only genes with at least a 0.25 log-fold change and expressed in at least 25% of the cells in the cluster under comparison were evaluated. Cluster similarity was assessed using the *BuildClusterTree* function of *Seurat*.

All violin plots, t-SNE visualisations and heatmaps were generated using functions from *Seurat, ggplot2, pheatmap, and grid* R packages. Construction of t-SNEs were achieved using the same number of principle components as the corresponding clustering and using default *Seurat* perplexity. DEGs from each cluster were used for pathway analysis using Bioreactome(Fabregat et al., 2018).

### Statistical analysis

No randomization and no blinding was used for the animal experiments. Animals were excluded from the omentectomy analysis if omentectomy was found to have been incomplete. Whenever possible, the investigator was partially blinded for assessing the outcome (bacterial binding). All data were analyzed using Prism 7 (GraphPad Prism, La Jolla, CA). Statistical tests performed for each data set are described within the relevant figure legend.

### Data availability

The authors declare that all relevant data supporting the finding of this study are available on request. R scripts for performing the main steps of analysis are available from the corresponding authors on reasonable request. scRNA-seq data sets have been deposited at GEO, accession GSM4053741.

---

**Supplementary Figure 1: DEG distinguishing omentum stromal cell subsets**

Violin plots of canonical omental stromal cell gene expression by cluster with highest log-normalized expression value labelled. Plots highlighting DEGs characteristic of mesothelial (**A,F**) immune (**B,C**) fibroblast (**D**) and ECM (**E**).

**Supplementary Figure 2: Pathway analysis of omentum stromal cell subsets**

Bioreactome analysis of DEGs for each cluster.

**Supplementary Figure 3: Adipose tissue fibroblasts express CCL11 whereas FALC FRC and mesothelial cells do not**

Representative confocal image of naive whole mount immunofluorescence staining of omentum, DAPI (Blue), PDPN (Green), CD45 (Yellow), CCL11 (Pink). Upper panel showing omental adipose in the absence of FALC, lower panel CD45^+^ FALC within omental adipose. FALC boundary delineated by dotted line in lower pane. Scale Bar 50μM. Staining representative of n=4 mice in 2 independent experiments.

**Supplementary Figure 4: Flow-cytometric gating strategy of murine PEC and Omentum digests.**

PEC and Omentum were analysed by dead cell exclusion, determination of single cell populations (including B cells) using scatter profiles and based on CD45 positivity. CD19^+^ B-cells, TCRβ^+^ T cells and Siglec F^+^ Eosinophils were excluded, Ly6G^+^ neutrophils were gated.

**Supplementary Figure 5. Neutrophils do not aggregate on the parietal wall during peritonitis**

Representative confocal images of immunofluorescence staining of murine parietal wall biopsies taken from naïve or 12h post injection of Zymosan A. CD11b (Magenta) MPO (White). Scale bar 50μM.

**Supplementary Figure 6: CitH3+ DNA covers neutrophil aggregates forming on omFALCs during peritonitis.**

**A**, Confocal image showing inside the cluster presented in Fig. 4C obtained by wholemount immuno-fluorescence staining of omentum from mice 6 hours post-Zymosan injection showing DAPI (magenta), CitH3 (Green) and Ly6G (Blue). **B**, Confocal image showing inside another representative cluster 6 hours post-Zymosan injection (first panel) and 3D reconstruction of whole omFALC showing the surface of the cluster (first row) and a clipped view of the inside of the cluster (second row). Scale Bar 100μM.

**Supplementary Table 1: Summary table of patient demographics.**

Sex, Age, body mass index (BMI) white blood cell (WBC) count and C-reactive protein (CRP) characteristics of the Biliary colic patients and acute appendicitis patients recruited for the study.

**Supplementary Table 2: Antibodies used for the study**

Primary and secondary antibodies used in the murine and human studies.

**Supplementary Table 3: List of differentially expressed genes**

Differentially expressed genes were identified as genes with a 0.25 log-fold change and expressed in at least 25% of the cells in the cluster under comparison for each cluster

