## Supplementary Figures 1-6 & Table 1 & 2 for "Neutrophils mediate the capture of peritoneal contaminants by fat-associated lymphoid clusters of the omentum"

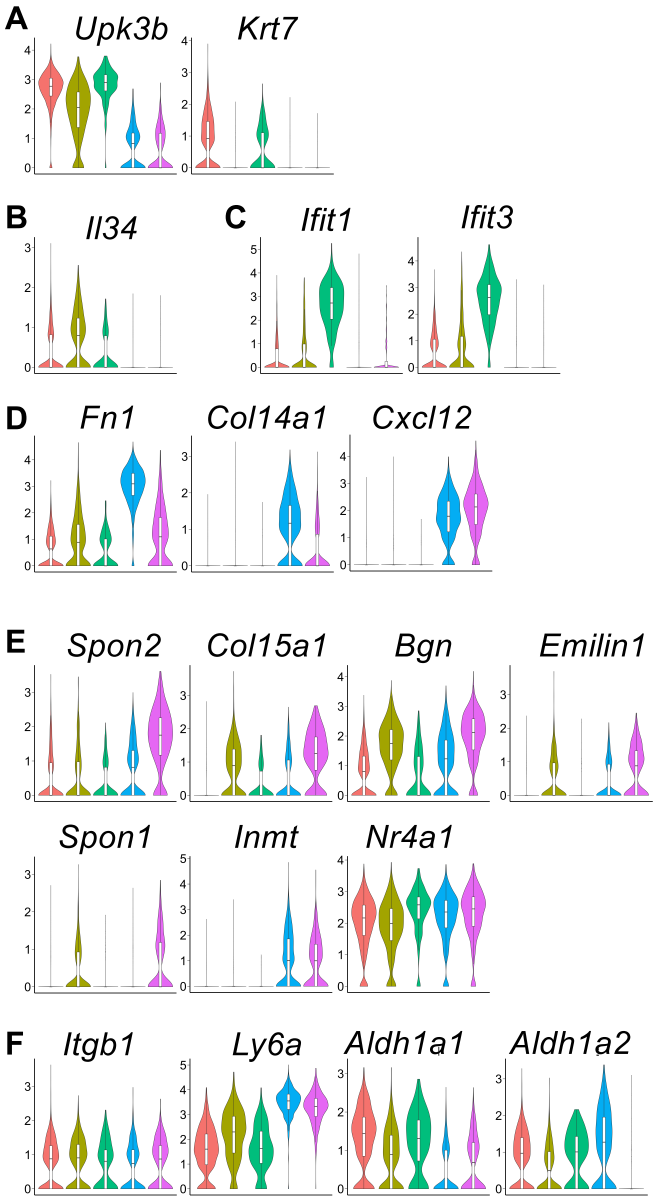


**Supplementary Fig. 1**

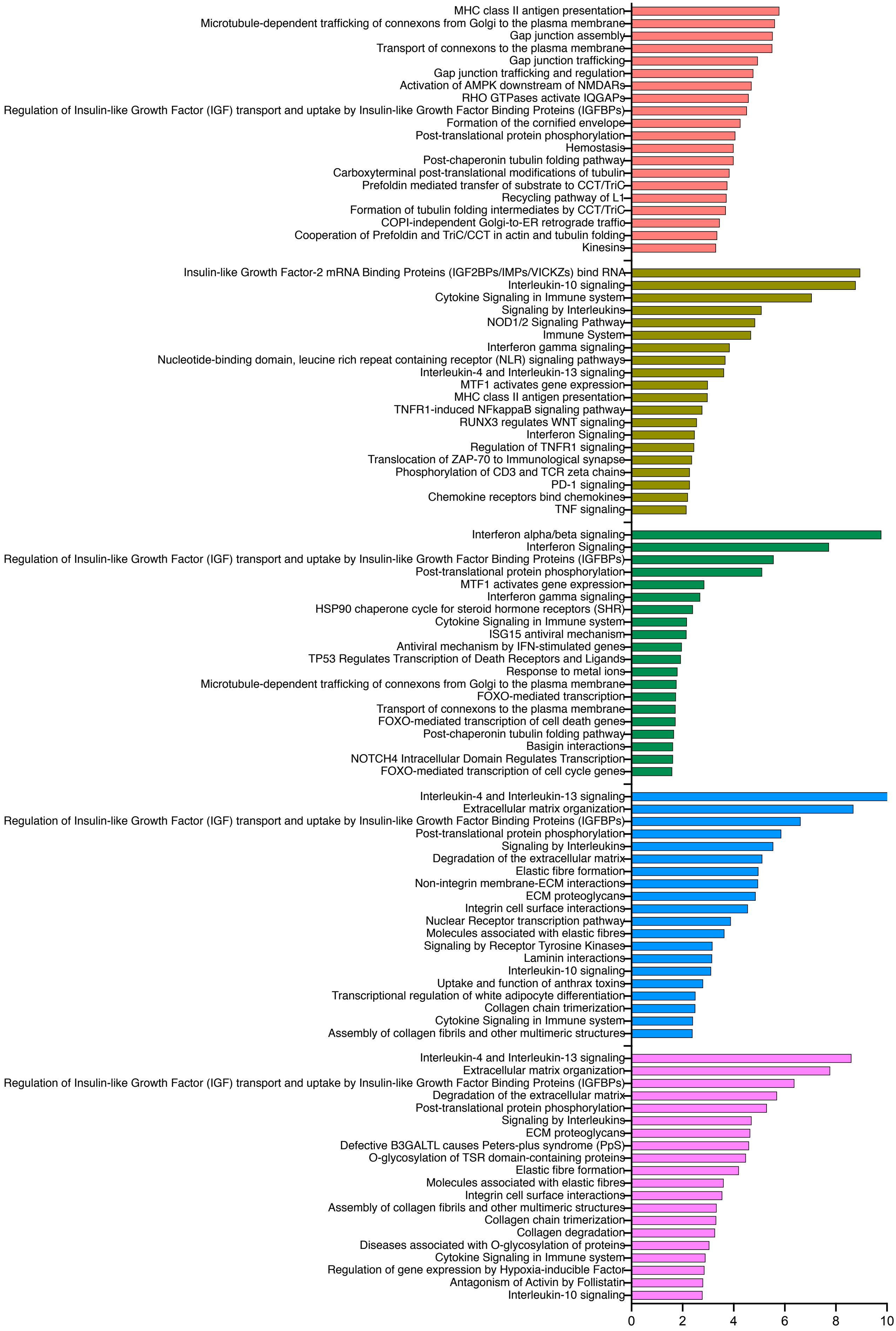


**Supplementary Fig. 2**

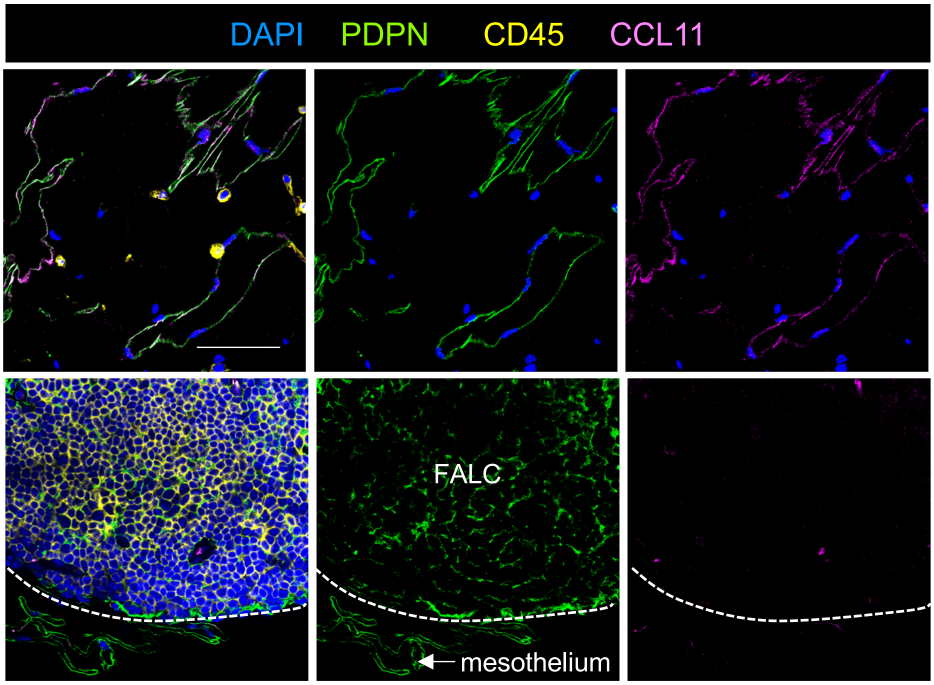


**Supplementary Fig. 3**

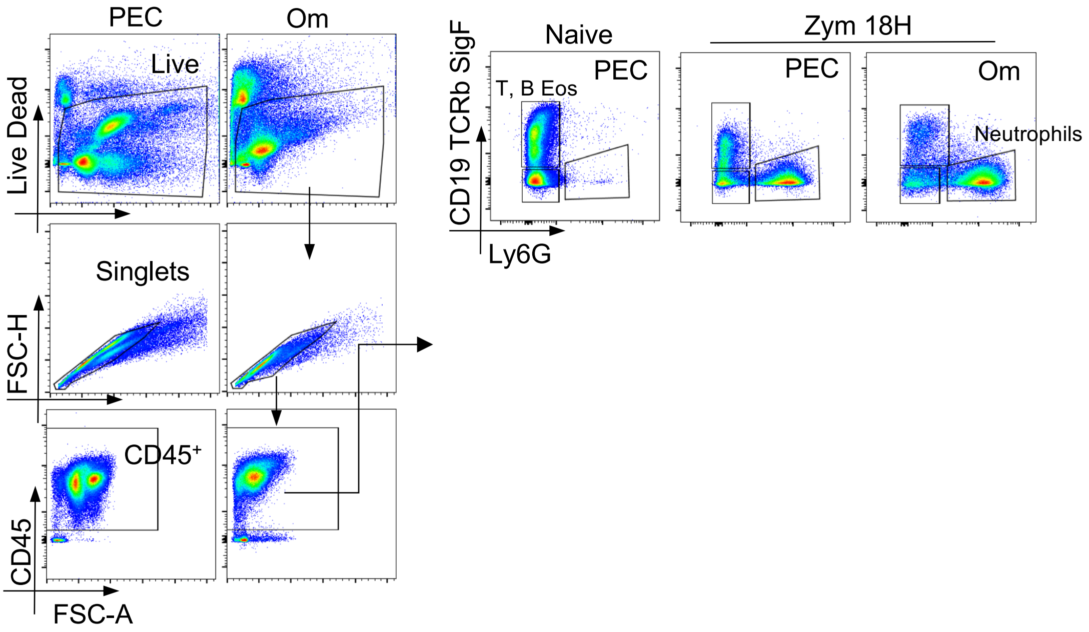


**Supplementary Fig.4**

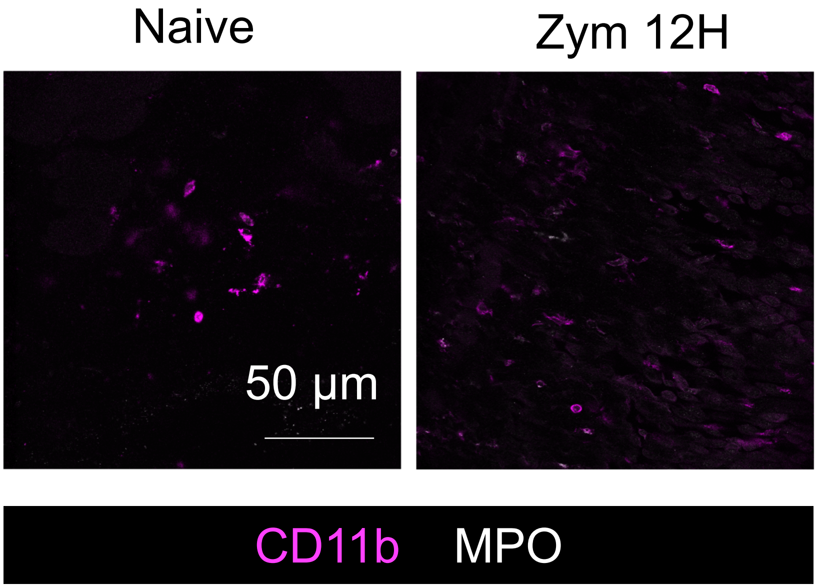


**Supplementary Fig.5**

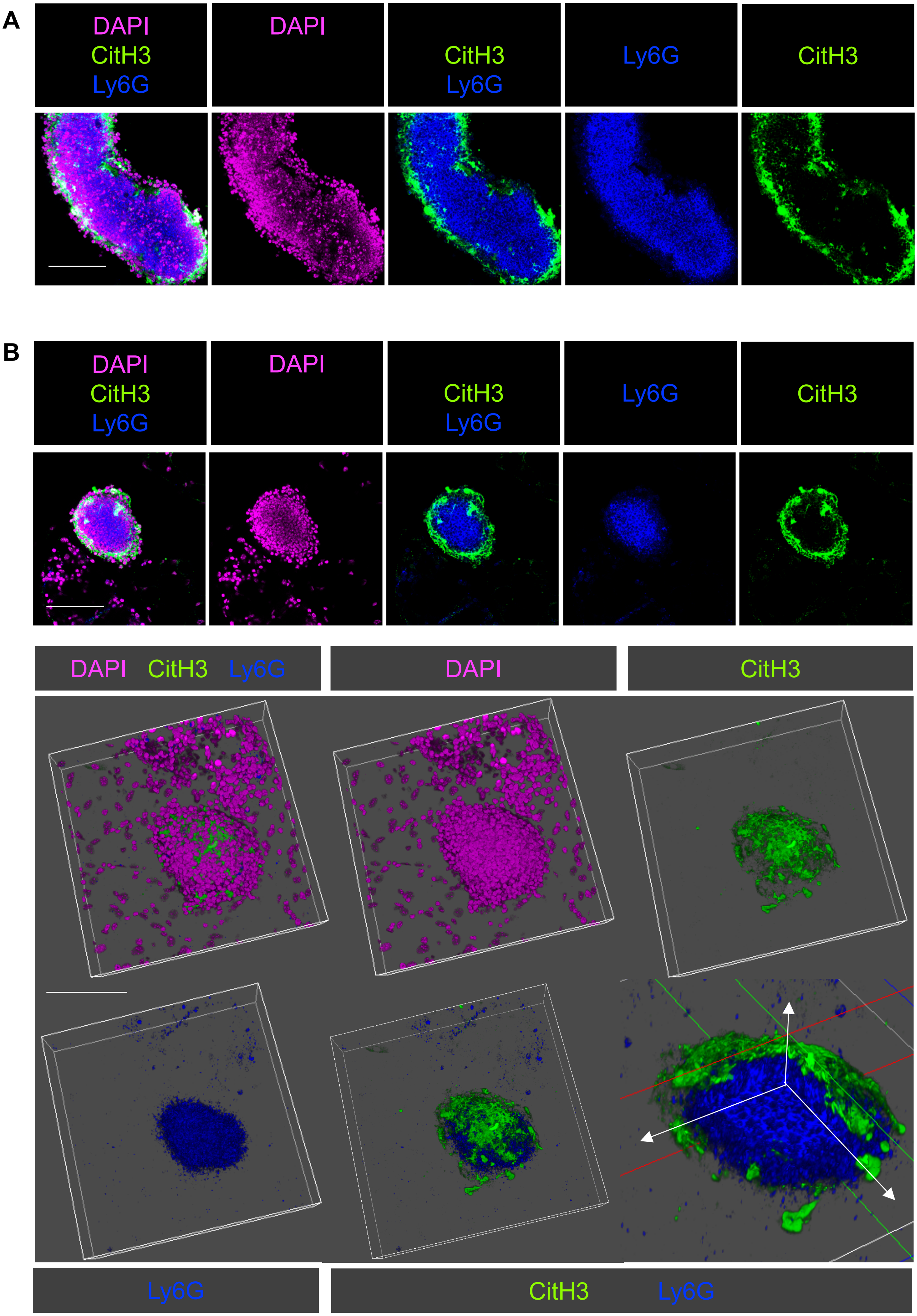


**Supplementary Fig. 6**

| | Acute appendicitis | Biliary colic |
|---|---|---|
| **Number of participants** | 13 | 10 |
| **Female:Male** | 7:6 | 8:2 |
| **Median age in years (range)** | 28 (16 – 58) | 48 (22 to 67) |
| **Body mass index (mean)** | 29.1 | 32.3 |
| **White blood cell count x 10^9/L*** | 14.78 (5.2 - 19.7) | 6.81 (3.90 - 9.80) |
| **Neutrophil count x 10^9/L*** | 11.95 (3.16 - 16.81) | 4.05 (2.16 - 8.08) |
| **Lymphocyte count x 10^9/L*** | 1.63 (0.38 - 2.90) | 1.99 (1.13 - 2.74) |
| **Monocyte count x 10^9/L*** | 1.10 (0.14 - 2.06) | 0.54 (0.35 - 0.75) |
| **Eosinophil count x 10^9/L*** | 0.07 (0.01 - 0.06) | 0.24 (0.05 - 62.00) |
| **Basophil count x 10^9/L*** | 0.02 (0.01 - 0.06) | 0.03 (0.01 - 0.05) |
| **Serum C-reactive protein mg/L** | 63 (6 to 215) | 4 (< 1 to 5) |

*data are mean (range)

**Supplementary Table 1**

| Reagent | Clone | Conjugate | Source |
|---|---|---|---|
| **Mouse primaries** | | | |
| Armenian Hamster anti-murine TCRb | h57-597 | BV421 | Biolegend |
| Donkey anti-mouse IgM | Polyclonal | Rhodamine Red | Jackson Laboratories |
| Goat anti-Cathepsin C | Polyclonal | Unconjugated | R&D Systems |
| Goat anti-CCL11/Eotaxin | Polyclonal | Unconjugated | R&D Systems |
| Goat anti-ENPP2/Autotaxin | Polyclonal | Unconjugated | R&D Systems |
| Goat anti-MPO | Polyclonal | Unconjugated | R&D Systems |
| Rabbit anti-CXCL1 | Polyclonal | Unconjugated | Abcam |
| Rabbit anti-histone H3 (citrulline R2 + R8 + R17) | Polyclonal | Unconjugated | Abcam |
| Rabbit anti-ISG15 | Polyclonal | Unconjugated | ThermoFisher |
| Rat anti-I-A/I-E | M5/114.15.2 | AF780 | eBiosciences |
| Rat anti-CD11b | M1/70 | PE/Dazzle594 | Biolegend |
| Rat anti-CD19 | 6D5 | BV421 | Biolegend |
| Rat anti-CD31 | 390 | Pacific blue | Biolegend |
| Rat anti-CD41 | MWReg30 | APC-Cy7 | Biolegend |
| Rat anti-CD44 | IM7.8.1 | PE | Miltenyi |
| Rat anti-CD45 | 104 | BV650<br>APC-Cy7 | Biolegend |
| Rat anti-CD55 | REA300 | FITC | Miltenyi |
| Rat anti-CD200 | OX2 | PE | Biolegend |
| Rat anti-CXCL13 | DS8CX13 | APC | ThermoFisher |
| Rat anti-F4/80 | BM8 | PE-Cy7 | eBiosciences |
| Rat anti-Ly6C | HK1.4 | AF700 | Biolegend |
| Rat anti-Ly6G | 1A8 | BV421 | Biolegend |
| | | AF647 | Biolegend |
| Rat anti-Ly6G (Gr1) | RB6-8C5 | Biotin | eBiosciences |
| Rat-anti-CD140a | APA5 | PE | Biolegend |
| Rat anti-Siglec-F | E50-2440 | BV421 | BD Pharmingen |
| Rat anti-Ter119 | TER-119 | APC-Cy7 | Biolegend |
| Syrian hamster anti-Gp38 | 8.1.1 | APC | Biolegend |
| | | Pe-Cy7 | Biolegend |
| **Human primaries** | | | |
| Mouse anti-human CD3 | HIT3a | PE | Biolegend |
| | OKT3 | APC/Cy7 | Biolegend |
| Mouse anti-CD14 | HCD14 | Pacific Blue | Biolegend |
| Mouse anti-CD15 | W6D3 | PE/Dazzle 594 | Bioleged |
| Mouse anti-CD19 | HIB19 | PE | Biolegend |
| Mouse anti-CD19 | HIB19 | PE/Cy7 | Biolegend |
| Mouse anti-CD45 | HI30 | Brilliant Violet 650 | Biolegend |
| Mouse anti-CD56 | MEM-188 | PE | Biolegend |
| Mouse anti-CD64 | 10.1 | APC/Cy7 | Biolegend |
| Mouse anti-CD163 | GHI/61 | Brilliant Violet 605 | Biolegend |
| Mouse anti-CD206 | 15--2 | Alexa Fluor 647 | Biolegend |
| Mouse anti-HLA-DR | L243 | FITC | Biolegend |
| Mouse anti-human IgM | MHM-88 | Alexa Fluor 647 | Biolegend |
| Rat anti-CD11b | M1/70 | Alexa Fluor 700 | Biolegend |
| **Secondaries** | | | |
| Donkey anti-Goat | Polyclonal | AlexaFluor488 | Invitrogen |
| Donkey anti-Rabbit | Polyclonal | AlexaFluor555 | Invitrogen |
| Streptavidin | | AlexaFluor555 | Invitrogen |
| | | PerCP | Biolegend |
| | | APC | Biolegend |
| | | BV711 | Biolegend |

**Supplementary Table 2**
